# Saturated reconstruction of living brain tissue

**DOI:** 10.1101/2022.03.16.484431

**Authors:** Philipp Velicky, Eder Miguel, Julia M. Michalska, Donglai Wei, Zudi Lin, Jake F. Watson, Jakob Troidl, Johanna Beyer, Yoav Ben-Simon, Christoph Sommer, Wiebke Jahr, Alban Cenameri, Johannes Broichhagen, Seth G. N. Grant, Peter Jonas, Gaia Novarino, Hanspeter Pfister, Bernd Bickel, Johann G. Danzl

## Abstract

Complex wiring between neurons underlies the information-processing network enabling all brain functions, including cognition and memory. For understanding how the network is structured, processes information, and changes over time, comprehensive visualization of the architecture of living brain tissue with its cellular and molecular components would open up major opportunities. However, electron microscopy (EM) provides nanometre-scale resolution required for full *in-silico* reconstruction^1–5^, yet is limited to fixed specimens and static representations. Light microscopy allows live observation, with super-resolution approaches^6–12^ facilitating nanoscale visualization, but comprehensive 3D-reconstruction of living brain tissue has been hindered by tissue photo-burden, photobleaching, insufficient 3D-resolution, and inadequate signal-to-noise ratio (SNR). Here we demonstrate saturated reconstruction of living brain tissue. We developed an integrated imaging and analysis technology, adapting stimulated emission depletion (STED) microscopy^6,13^ in extracellularly labelled tissue^14^ for high SNR and near-isotropic resolution. Centrally, a two-stage deep-learning approach leveraged previously obtained information on sample structure to drastically reduce photo-burden and enable automated volumetric reconstruction down to single synapse level. Live reconstruction provides unbiased analysis of tissue architecture across time in relation to functional activity and targeted activation, and contextual understanding of molecular labelling. This adoptable technology will facilitate novel insights into the dynamic functional architecture of living brain tissue.

## Introduction

Brain computation and information storage are intimately linked to the structure of a network of ∼86 billion neurons^15^ in humans. Each is typically connected by thousands of information-transmitting synapses to other neurons and interacts with glial support cells. To address how this incredibly crowded and complex tissue’s architecture, connectivity, and functional activity evolve over time and interrelate, one would ideally employ a technology that enables imaging and *in-silico* reconstruction of *living* brain tissue. This would allow mapping of how neuronal and non-neuronal cells and their delicate, functionally paramount subcellular specializations, such as synapses, relate to each other in the 3D tissue environment, and how this changes over time or in response to specific intervention. Combining connectivity information with the location of specific molecules could then further define cellular and subcellular identities, and provide a molecular ground truth for synapse location and type.

EM reconstruction^1–5^ offers the most detailed insights into brain architecture. However, it is limited to fixed specimens, static tissue representations, and sample preparation that impedes molecular labelling. A light microscopy-based technology for tissue reconstruction, in contrast, could enable observation of living systems, together with visualization of specific molecules and cellular signaling. The intricate cellular arrangements in brain tissue call for a super-resolution approach^6–9^ with resolution improvement in all three spatial dimensions, because reconstruction is limited by the lowest-resolution direction (typically along the optical axis, *z*-direction). Conventional (diffraction-limited) microscopy is unsuitable, as it has a best-case lateral resolution of about half the wavelength of employed light and axial resolution as poor as ∼1000 nm for tissue-compatible high-numerical aperture objective lenses and far red excitation. The ongoing revolution in machine learning is transforming not only image analysis, including for connectomics, but brings about new concepts for how to amalgamate data acquisition and analysis in microscopy.

Here we introduce saturated reconstruction of living brain tissue. To achieve dense reconstruction of the cellular components in tissue volumes, we developed an integrated optical /machine learning technology for tissue imaging that breaks the intertwined limitations for 3D-resolving power, signal-to-noise ratio (SNR), speed, and light burden in classical super-resolution imaging of living systems. We started from the realization that this would require specifically adapting and melding the three key elements factoring into overall information yield, i.e. biological preparation, in-tissue 3D-super-resolution imaging technology, and analysis. We based our technology on stimulated emission depletion^6,13^ (STED) microscopy. Here, a light pattern turns off fluorophores at sub-nanosecond timescales except those located near its intensity minimum and positions are queried sequentially. STED is thus compatible with freely diffusing fluorophores and relatively robust against movement in living samples. Unlike visualization of protein distributions or tracing the morphology of sparse cells^16^, saturated tissue reconstruction requires comprehensive delineation of all cells. We therefore employed super-resolution shadow imaging^14^, where extracellularly applied fluorophores^17^ define cellular structures more comprehensively than in single-molecule approaches for extracellular space imaging (ECS)^18^, and which has proven powerful for visualizing cellular arrangements and neuronal interactions^19–21^. Here, photobleached fluorophores are replenished by diffusion, and radicals generated from extracellular fluorophore bleaching are less able to damage the specimen than intracellular radicals. Despite these advantages, synapse-level reconstruction of living brain tissue has been elusive. The square-root dependence of resolution on applied STED laser power^22^ and the need for Nyquist sampling in 3D with low tens of nanometers step sizes impose a heavy cost of tissue light-burden to increase 3D-resolution^23^. Together with in-tissue optical imperfections leading to progressive signal loss at higher resolution, these factors ultimately limit achievable 3D-resolution and signal-to-noise ratio (SNR)^24^.

We therefore specifically modified STED for improved SNR and near isotropically super-resolved tissue imaging, coupled with a custom-designed, two-stage deep-learning strategy. Stage one leveraged information on sample structure from numerous separate, prior measurements allowing to drastically reduce light burden and imaging time without sacrificing resolution, and hence enabled live-tissue compatible volumetric super-resolution imaging. Counterintuitively, SNR was improved even beyond ground-truth high-SNR training data, which was key for subsequent automated segmentation tasks. Stage two was adapted from EM-connectomics algorithms and trained to automatically translate our volumetric live-imaging data into a saturated instance segmentation at single synapse level. We termed this technology LIONESS for *Live Information-Optimized Nanoscopy Enabling Saturated Segmentation* (**Fig. 1A**). LIONESS allows dynamic brain tissue reconstruction paired with molecular and functional information.

**Fig. 1.**
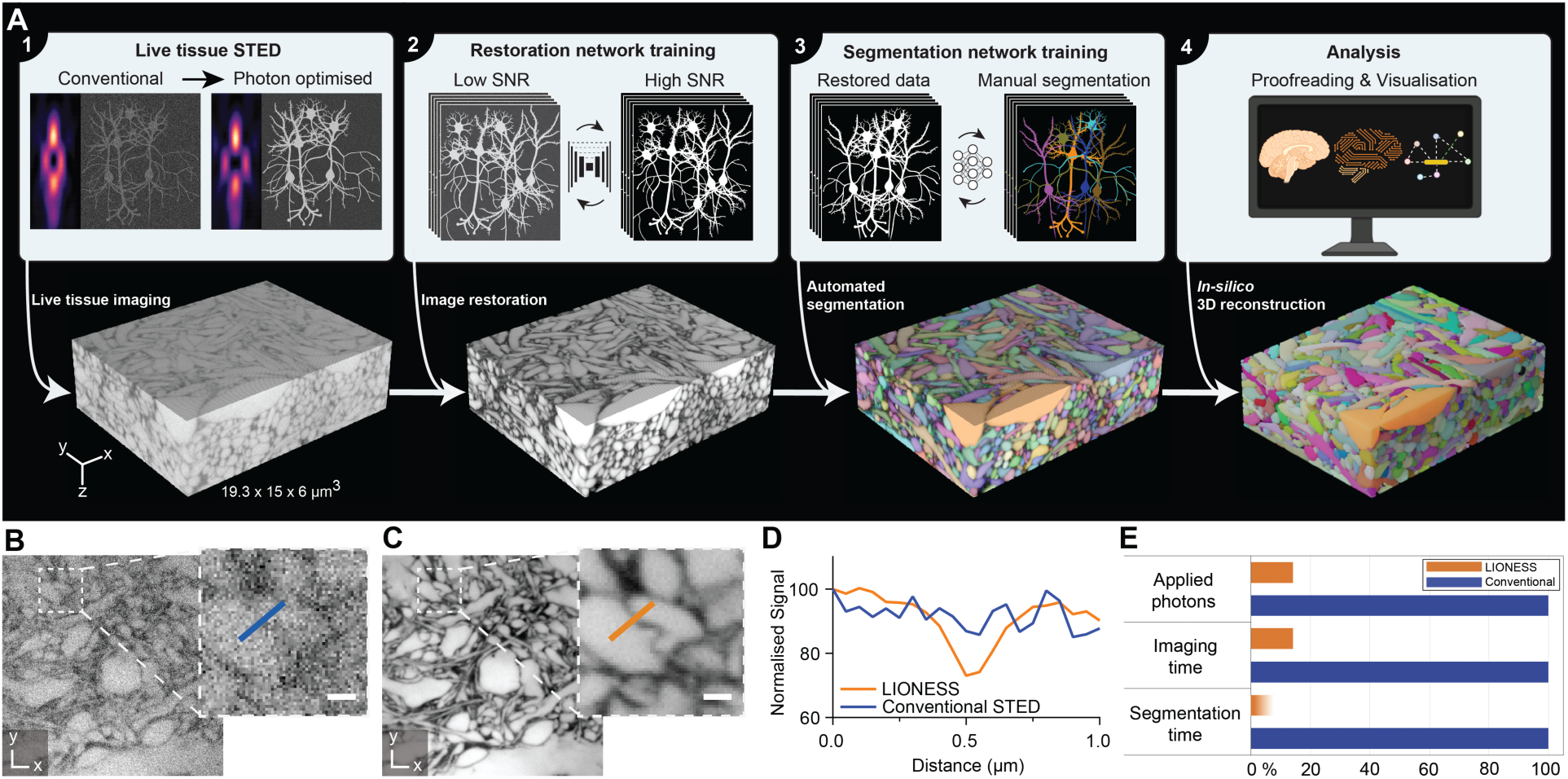
LIONESS enables saturated reconstruction of living brain tissue. (**A**) LIONESS technology exemplified in living human cerebral organoid. Optical improvements, deep-learning training, and analysis (top) flow into individual processing steps (bottom): (1) Near-infrared STED with light patterns for improved effective point-spread-function in tissue. (2) Deep neural network training on paired low-exposure, low-SNR and high-exposure, high-SNR 3D-super-resolved volumes. (3) Deep 3D-segmentation network training with manually annotated data. (4) Postprocessing. (**B**) Conventional STED imaging in CA1 neuropil of organotypic hippocampal slice culture with phase modulation patterns for lateral (*xy*)- plus axial (*z*)-resolution increase. (**C**) Same region imaged with tissue-adapted STED patterns (4π-helical plus π-top-hat phase modulation), improved detector dynamic range, and deep-learning based SNR-restoration. STED power and dwell time were identical in panels B and C. Scale bars: 500 nm. (**D**) Line profiles across a synaptic cleft as indicated in (B) and (C). (**E**) Light exposure and imaging time were reduced by 86% with LIONESS compared to conventional high-photon load ground truth in network training, in addition to optical improvements of SNR. Deep-learning accelerates segmentation by orders of magnitude, also depending on sample complexity. LIONESS lookup tables are linear and inverted throughout, ranging from black (maximum photon counts extracellularly) to white.

## Results

### Achieving near-isotropic high-SNR STED in tissue

We opted for near-infrared STED (775 nm), to deliver highest STED performance coupled with reduced tissue absorption and scattering over visible light^23,25^. We screened for cell-impermeant fluorophores to label ECS and delineate cellular structures with maximized extra- vs. intracellular contrast, and identified suitable hydrophilic, anionic high-performance STED labels, including both unmodified commercial and custom sulfonated variants (**Suppl. Fig. 1**). We aimed for near-isotropic STED resolution and first incoherently overlapped classical 2π-helical and π-top-hat phase modulation patterns to achieve lateral (*xy*) and predominantly axial (*z*) STED resolution increase^24^, respectively, and mitigated spherical aberrations on the sensitive *z*-STED pattern using a silicone immersion objective with correction collar. However, the resulting intensity minimum of the combined light patterns was highly susceptible to aberrations and imperfect spatial overlap in tissue. We therefore replaced the 2π(*xy*)-pattern with a helicity-2 mode generated by 4π-helical phase modulation^26^. The shallower intensity rise and broader overall distribution of the 4π(*xy*)-STED pattern allowed more robust in-tissue alignment and improved quenching of “sidelobe” fluorescence insufficiently silenced by the *z*-STED pattern alone (**Suppl. Fig. 2A-C**). This led to substantially enhanced definition of cellular structures (**Suppl. Fig. 2D**). Increasing detector dynamic range further improved SNR (**Suppl. Fig. 3**). To delineate narrow spaces between cells with the extracellular label in 3D and detect the fluorescence modulation produced by thin cellular processes with sufficient SNR for segmentation, we integrated photons for 70µs per 50 x 50 x 50 nm^3^ voxel and dialed in near-isotropic resolution of ∼130 nm. However, this approach was too harsh and too slow for volumetric imaging of living tissue, causing dramatic photodamage in the tissue sample (**Suppl. Fig. 4A)**.

### Low-exposure, high-speed STED

We thus sought for strategies to simultaneously reduce both the light burden and imaging time while augmenting SNR. To achieve this, we recorded low-exposure STED data at high speed and deployed deep-learning-based restoration to computationally increase SNR, retrieving information on sample structure from prior measurements. We trained a convolutional neural network^27^ (**Suppl. Fig. 5A)** on paired low- and high-SNR imaging volumes from mouse organotypic slice cultures and the alveus region of acutely prepared mouse hippocampus. These contained diverse structures that were sampled at high SNR with 70 µs voxel dwell time, from which we set aside photon counts of the first 10 µs as low-SNR training input data. This ensured that both corresponded to voxel-exact equivalent sample structures. We could then apply this trained neural network to previously unseen data to predict high-SNR images from low-exposure input data. To evaluate the accuracy of this prediction in the context of cellular segmentation, per-voxel probabilistic uncertainty measures and ensemble disagreement between independently trained networks^27^ were of limited utility (**Suppl. Fig. 5B**). Therefore, we compared prediction outcome with corresponding sparse, positively labelled cellular structures (**Suppl. Fig. 6A**) and with paired high-SNR measurements, with both not included in the training (**Suppl. Fig. 6B**). This indicated that inaccuracies at the voxel level did not negatively impact delineation of cellular structures.

Repeated volumetric imaging in this low-exposure mode left cells intact, whereas they disintegrated when aiming to achieve similar resolution and SNR with the conventional high-photon load STED mode (**Suppl**. **Fig. 4**). Development of this scheme further reduced photon load by 86%. In contrast to current techniques^24^ for reducing STED exposure^28,29^ and photobleaching^30,31^, it also correspondingly sped up acquisition 7-fold and additionally denoised the data. Integrating labelling, optimizations for in-tissue isotropically resolving super-resolution imaging, low-exposure data collection, and computational SNR restoration resulted in a substantial gain in image quality over conventional STED imaging for given live-tissue compatible STED light exposure (**Fig. 1B-E**). For example, when applied to dense neuropil in organotypic hippocampal slice cultures, this LIONESS imaging regime revealed synaptic clefts at STED light exposure far too low to do so in conventional STED mode (**Fig. 1B-D**). Together, this resulted in volumetric light-microscopy data suitable for comprehensive segmentation of living neuronal tissue.

### In-silico saturated reconstruction

Manual annotation of all cellular structures in a small volume of such LIONESS imaging data showed that saturated live tissue reconstruction was feasible. However, it was time consuming and therefore poorly scalable. Segmentation of a ∼400 µm^3^ volume of living brain tissue, selected from a highly interwoven region of neuropil in an organotypic hippocampal slice, took a trained segmenter ∼450 hours **(Suppl. Fig. 7)**. We therefore implemented a second deep neural network for automated segmentation, adapting algorithms and software from EM-connectomics^32,33^, and employed an iterative training scheme. We initially trained the network on a subvolume of the manually annotated LIONESS imaging volume (∼285 µm^3^, the other part was used for validation) and applied it together with watershed postprocessing to larger imaging volumes that harbored additional structural diversity (CA1 and dentate gyrus (DG) neuropil in organotypic hippocampal slice cultures and alveus in acutely prepared hippocampi). We then manually proofread the output and fed it back into training, thus extending the training volume to ∼800 µm^3^, yielding an improved segmentation neural network with enhanced prediction quality.

We chose living human cerebral organoids^34^ as first specimen for automated reconstruction, as these have emerged as powerful model systems for studying human brain development and disease mechanisms at cell to tissue level. When applied to a living cerebral organoid, our pipeline enabled saturated tissue reconstruction (**Fig. 1A**). Such samples with moderately complex tissue structure required minor intervention by manual proofreading. Reconstruction yielded contextual information not available with imaging of sparsely labelled specimens. For example, we observed the interaction of an axonal growth cone with neighboring structures in the living organoid (**Suppl. Fig. 8**). The gain in throughput from automated over manual segmentation was substantial, with our whole pipeline including microscopic data acquisition (140 seconds), image restoration (10 seconds) and automated saturated segmentation (∼40 minutes) taking less than 45 minutes excluding data inspection and proofreading (**Fig. 1E**). Manual segmentation would require an estimated ∼860 person-hours for this organoid dataset (1,737 µm^3^). Extracting the space not occupied by cellular segments additionally allowed us to reconstruct the ECS, which amounted to 225 µm^3^ or 13% in this organoid subvolume (**Suppl. Fig. 9**).

In the next step, we chose the alveus of intact, acutely dissected mouse hippocampi, a region that is extremely densely packed with thin neuronal processes transmitting signals to other regions. Automated reconstruction highlighted the thin, individually resolved axons running in various orientations and interacting with glial cells (**Suppl. Fig. 10A-C and Suppl. Videos 1 and 2**). Such dense but structurally comparatively homogeneous regions also required little intervention during proofreading. Approximately 45 corrections per mm axon length were necessary in such data, with false splits being the dominant type of error (**Suppl. Fig. 10D)**. These data showed that a comprehensive structural segmentation of living nervous tissue is feasible. Furthermore, deep-learning models for segmentation were applicable across brain tissue preparations.

### Validation of segmentation

To test the potential of this technology for saturated reconstruction and analysis of complex nervous tissue specimens, we collected imaging volumes from the highly interwoven neuropil in organotypic hippocampal slices. These volumes contained diverse neuronal structures from spiny dendrites, to axons and their boutons, and astrocytic processes. We focused on assignment of dendritic spines to dendrite branches, as the fine connecting spine necks are among the thinnest of neuronal structures. We tested our reconstruction capability against sparse structural ground truth from cytosolic EGFP expression, which revealed all dendritic spines on a given dendrite. From LIONESS imaging data alone, without automated segmentation, a neuroscientist blinded to EGFP ground truth data correctly assigned 73% (±8.3%, mean ± standard deviation, s.d.) of spines in four different example dendrite stretches (from three independent biological replicates; 129 spines in total, 34 missed, 2 false positive). When applied to the same datasets, the artificial network often segmented and correctly connected spines to the respective dendrite or classified spines as separate (orphan) segments which could then be unambiguously assigned to a dendrite. The experimenter who collected the data performed proofreading of the segmentation output and correctly attached 83% (±8.0%, mean ± s.d.; 129 spines in total, 20 missed, 0 false positive) of dendritic spines (**Fig. 2, Suppl. Fig. 11**). This showed that both manual and automated reconstruction can retrieve a large percentage of dendritic spines and demonstrated applicability of LIONESS for analysis of neuropil architecture.

**Fig. 2.**
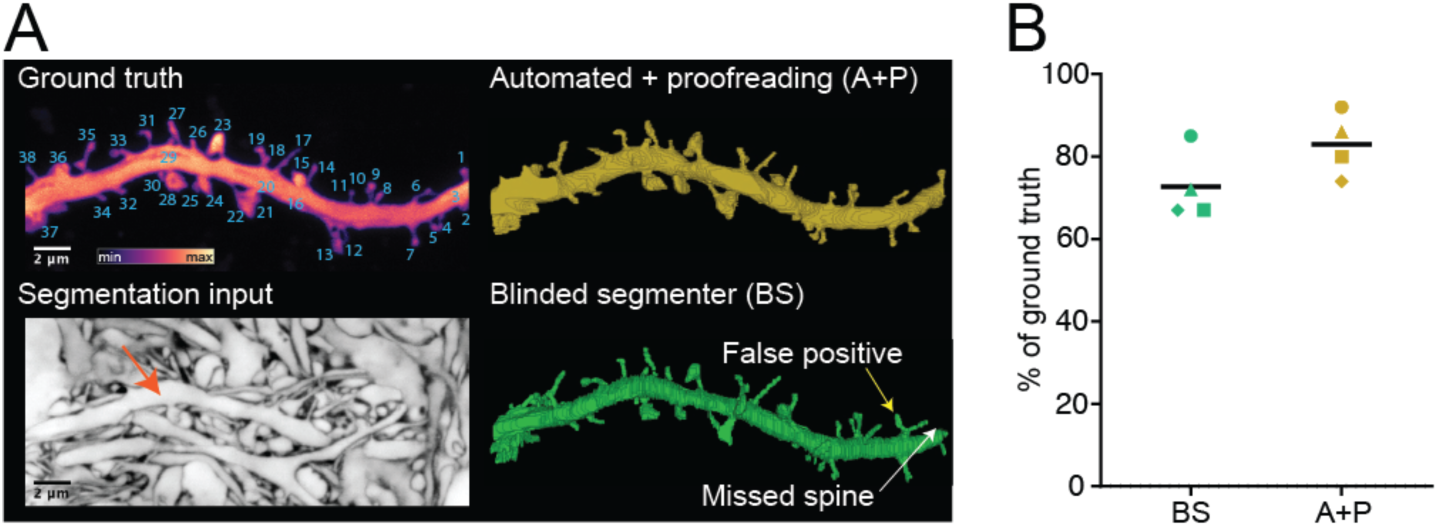
Validation of segmentation. **(A)** *Top left*: Maximum intensity projection of a dendrite in organotypic hippocampal slice culture labelled by cytosolic EGFP expression. The positive label serves as ground truth for segmentation, with individual spines numbered. *Bottom left*: Plane from volumetric LIONESS acquisition with arrow indicating the same dendrite (maximum intensity projection spanning 150 nm). *Top right*: 3D-reconstruction after automated segmentation and proofreading of LIONESS imaging data by the experimenter who recorded the data (A+P). As this person was not blinded to the EGFP-channel, this indicates which spines can be retrieved from LIONESS but does not serve as independent control. *Bottom right*: Fully manual spine detection from LIONESS imaging data by a segmenter blinded to the EGFP-channel (BS). Exemplary missed and false positive spines are indicated by white and yellow arrows, respectively. (**B)** Percentage of correctly assigned spines from the automated plus proofread (A+P, orange) and manual (BS, green) segmentations relative to the total number of spines in the positively labelled ground truth for 4 different datasets from three independent samples **(Suppl. Fig. 11)**. Correctly identified spines were counted as +1, false positives as -1. Black bars: mean of the 4 datasets.

### Connectivity reconstruction

We now applied our technology to living hippocampal neuropil in the dentate gyrus (DG) to unbiasedly visualize the architecture of this complex region. We identified and reconstructed diverse cellular constituents like myelinated and unmyelinated axons, spiny dendrites, and glial cells in the densely packed tissue volume (**Fig. 3A,B, Suppl. Fig. 12A, and Suppl. Video 3**). Similar as with EM, proofreading of automated segmentation remains a time-limiting factor, such that it is often preferable to selectively apply it to the specific structures of interest. We focused reconstruction on an individual dendrite, revealing 38 spines that showed various morphologies, including thin, branched, mushroom-shaped, and filopodial (**Fig. 3C**). Spine heads were of diverse 3D-shapes, some hand-like engulfing part of the presynaptic bouton. Spine lengths ranged from 0.54 µm to 3.96 µm (1.77 µm ± 0.69 µm, mean ± s.d.) and showed a unimodal distribution (**Fig. 3D**). We identified 28 axons where a bouton directly contacted a spine head, resulting in a total of 39 putative synapses along these 22 µm of reconstructed dendrite (**Fig. 3C, Suppl. Fig 12B)**. Most axons made single (20) or double (6) connections, however triple and quadruple spine contacting axons were also observed. We did not observe a preferred angular orientation of contacting axons with respect to the dendritic shaft, further demonstrating the complex arrangement of the DG neuropil. The mean density of spines was 1.7/µm dendrite length. Both length and density quantifications are in-keeping with previous data^35^. **Fig. 3E** details spine length and position along the dendrite for each spine, together with exemplary volumetric renderings of pre- and postsynaptic structures.

**Fig. 3.**
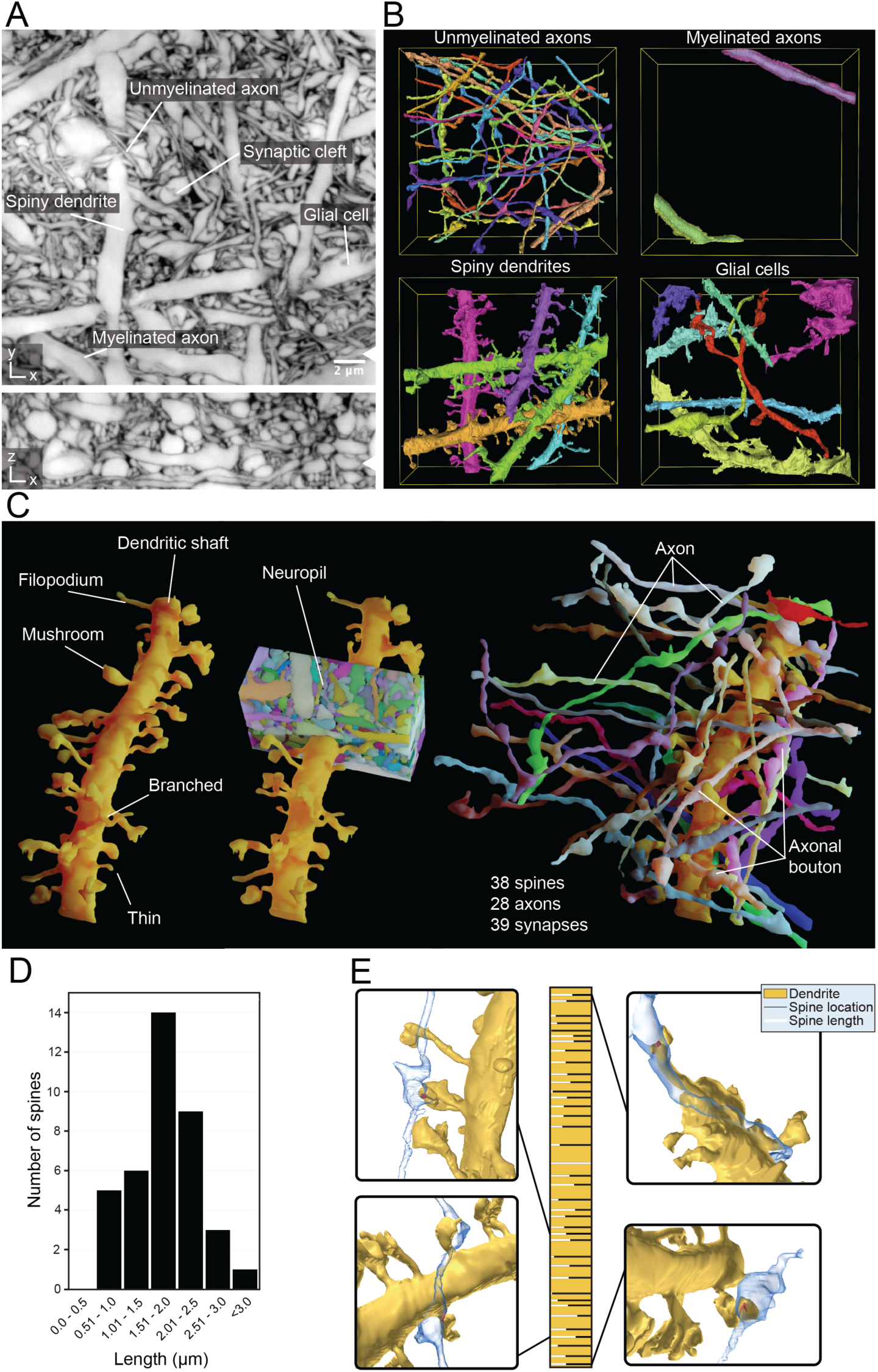
Connectivity reconstruction in live hippocampus. (**A**) Orthogonal planes from a LIONESS volume in *xy*- and *xz*-directions in neuropil of the dentate gyrus in organotypic hippocampal slice cultures. White arrowheads at image edges indicate position of corresponding orthogonal planes. Scale bar: 2 μm. LIONESS images are maximum intensity projections spanning 150 nm. (**B**) 3D-reconstructions of exemplary cellular structures extracted from (A). (**C**) 3D-reconstruction of a spiny dendrite from (A), showing various spine shapes (left), its embedding in dense neuropil (middle) and the 28 axons making a total of 39 synaptic connections at 38 spines (right). (**D**) Distribution of spine lengths for the dendrite in (C). (**E**) Spine location (bars) and relative spine lengths (white portion of bars) along the dendrite (gold) with 3D-renderings of exemplary synaptic connections.

### Molecularly informed reconstruction

We next sought to integrate key methods for live molecular labelling into tissue reconstruction. Live affinity labels proved useful for corroborating identity of specific structures, like myelinated axons (**Suppl. Fig. 13**). Most importantly, light microscopy is unrivalled at visualizing the location of specific proteins. As mere proximity of spines and boutons can be a poor predictor of synaptic connectivity between neurons^1^, we complemented it with a molecular definition of synaptic sites. We used a mouse line expressing a HaloTag fused to endogenous PSD95 protein^36,37^, an abundant protein located in the postsynaptic densities of excitatory synapses. Irreversible binding of applied HaloTag ligands coupled to a fluorescent dye visualized all excitatory postsynapses. In addition, we applied a combination of adeno-associated virus (AAV) and pseudotyped rabies particles^38^ to induce expression of EGFP-coupled synaptophysin, visualizing pre-synaptic terminals (**Fig. 4A, Suppl. Fig. 14, Suppl. Video 4**). These pre- and postsynaptic markers were combined with 3D-structural LIONESS imaging in the CA1 neuropil, providing cellular context lacking with conventional molecular labelling and imaging. Combined structural and molecular information unambiguously revealed various types of connections between pre- and postsynaptic partners: boutons of two separate axons converging on a single spine, single boutons contacting two neighboring spines of the same dendrite, and textbook-like single bouton to single spine connections (**Fig. 4B**). Excitatory synapses are preferentially located at dendritic spines, however also occur on dendritic shafts, in particular on aspiny interneurons. We also used the combined molecular and structural information to determine the fraction of excitatory synapses with dendritic shaft location in our imaging volumes, equaling 8.3% in **Fig. 4** and 14.7% in **Suppl. Fig. 14**. Of note, comparison with diffraction-limited readout of synaptic molecules further illustrated the gain in 3D-definition with LIONESS (**Fig. 4A**, **bottom**, **Suppl. Fig. 14B, bottom**).

**Fig. 4.**
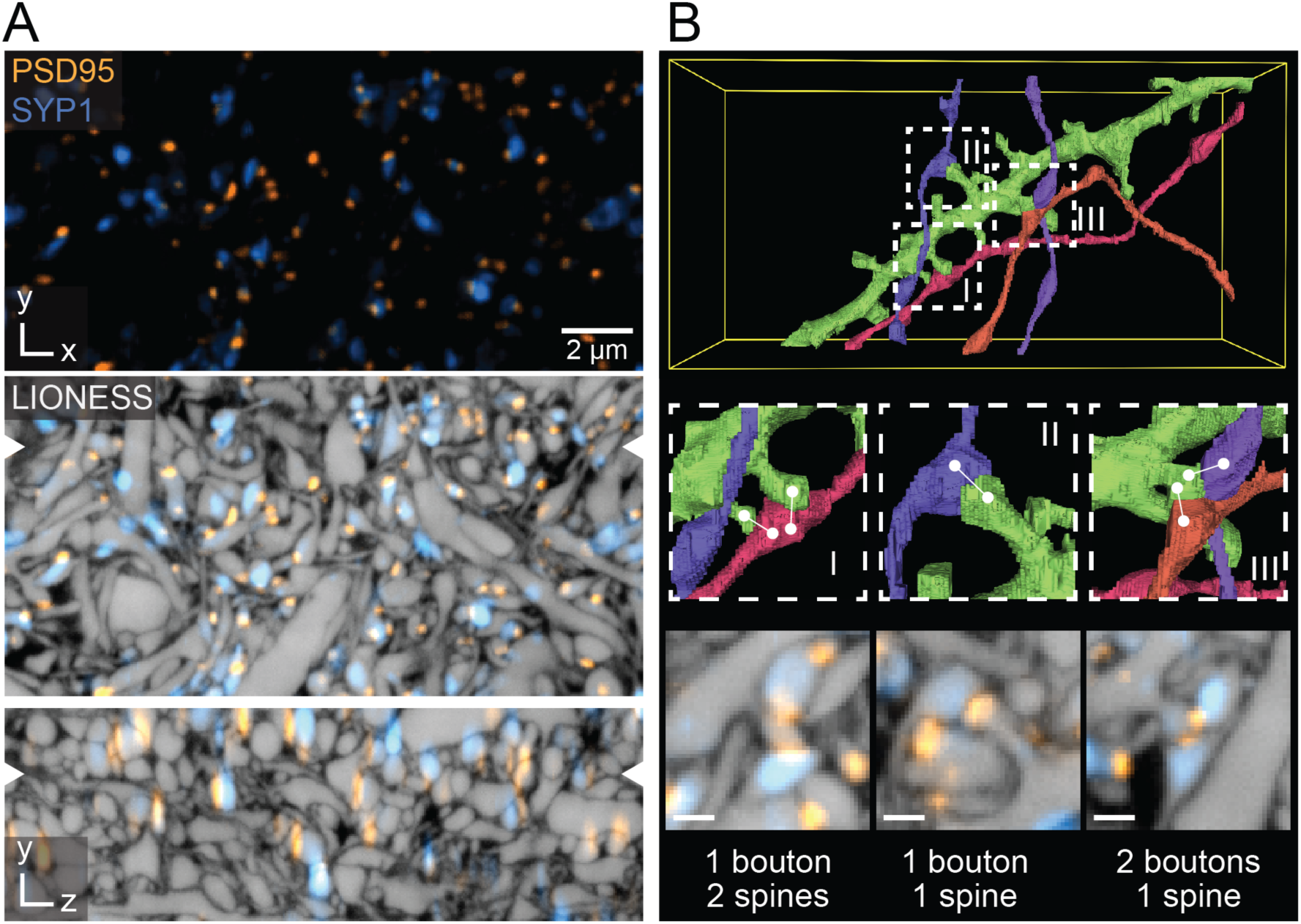
Molecularly informed reconstruction of living brain tissue. (**A**) *Top*: Confocal image of CA1 neuropil in organotypic hippocampal slice culture, with virus-assisted delivery of synaptophysin 1-EGFP (SYP1, blue) highlighting a subset of presynaptic terminals, and PSD95-HaloTag knock-in labelling all excitatory postsynapses. Denoising applied. Scale bar: 2 µm. *Middle* and *bottom*: Overlay with near-isotropically super-resolved volumetric LIONESS data. Orthogonal planes in *xy*- and *xz*-directions represented as maximum intensity projections spanning 150 nm, with positions of corresponding planes indicated by arrowheads at image edges. Diffraction-limited SYP1 and PSD95 signals extend beyond the cellular structures defined by LIONESS. (**B**) *Top*: 3D-reconstruction of a selected dendrite (green) from the same LIONESS volume with 4 synaptically connected axons. *Middle*: Magnified views as indicated by the dashed boxes in the top panel, highlighting diverse geometric arrangements of boutons and spines. White lines indicate synaptic connections retrieved from the molecular information. *Bottom*: LIONESS planes from the corresponding subvolumes together with molecular information. Maximum intensity projections spanning 150 nm. Scale bars: 500 nm.

### Morphodynamics and activity

Our low-exposure approach allowed repeated reconstruction of the same tissue volume, revealing how subcellular morphologies and the neuronal network evolved over time, while pairing this directly with optical readout of activity. We first used LIONESS to repeatedly image the same volume of hippocampal neuropil over 3 days. This allowed us to observe morphology changes and movement of neuronal and non-neuronal subcellular structures in their context (**Suppl. Fig. 15**), rather than being limited to sparse, positively labelled cells^16^.

We now devised an all-optical approach to correlate 3D-structure and signaling activity in the same living cellular network. We focused on the hippocampal circuitry, where mossy fibers originating from dentate gyrus (DG) granule cells deliver excitatory input to the proximal dendrites of pyramidal neurons in the CA3 region, forming boutons on complex spines often termed thorny excrescences^39^ (**Suppl. Fig. 16A,B**). Using organotypic slices from mice where all DG granule cells expressed the calcium indicator^40^ GCaMP6f (Ai95/Prox1-cre) we recorded calcium transients in individual mossy fiber boutons with sub-second resolution, applying pharmacological manipulation with the GABA_A_ receptor antagonist gabazine to enhance network activity. LIONESS revealed the underlying 3D-cellular organization (**Suppl. Fig. 16C,D, Suppl. Video 5, 6**). When we repeated volumetric LIONESS imaging, both mossy fiber boutons and their postsynaptic complex spines showed structural dynamics on the minutes timescale. Signaling activity continued during LIONESS acquisition (**Suppl. Fig. 16B,E**).

We next developed a more refined approach for investigating activity and dynamics within the tissue, combining chemogenetically targeted cell activation with Ca^2+^-imaging and dynamic reconstruction in the same living specimen. We expressed the virally encoded DREADD (designer receptor exclusively activated by designer drugs)^41^ hM3Dq in a subset of DG granule cells, which enhanced neuronal excitation upon application of the bio-orthogonal drug clozapine N-oxide (CNO). This allowed us to control and image the activity of a large mossy fiber bouton in the DG hilus, before structurally reconstructing it together with the complex spines on the postsynaptic hilar mossy cell (**Fig. 5A, Suppl. Video 7**). Comprehensive automated segmentation additionally clarified the relationship with neighboring mossy fiber boutons **(Suppl. Video 8)**. We visualized the structural evolution after 19.5h (**Fig. 5B**), which revealed dramatic rearrangements in synaptic architecture, mirrored in a bouton volume change from 11.8 µm^3^ to 8.3 µm^3^. These values are comparable to volumes of large mossy fiber boutons on CA3 pyramidal cells determined by serial sectioning EM in rat hippocampus^39^. However, with its applicability to living tissue, LIONESS has the unique capacity to repeatedly retrieve both activity and dynamic structural information directly in the living state. It has thus the capability to follow structural plasticity and determine structure-function relationships in neuronal tissue.

**Fig. 5.**
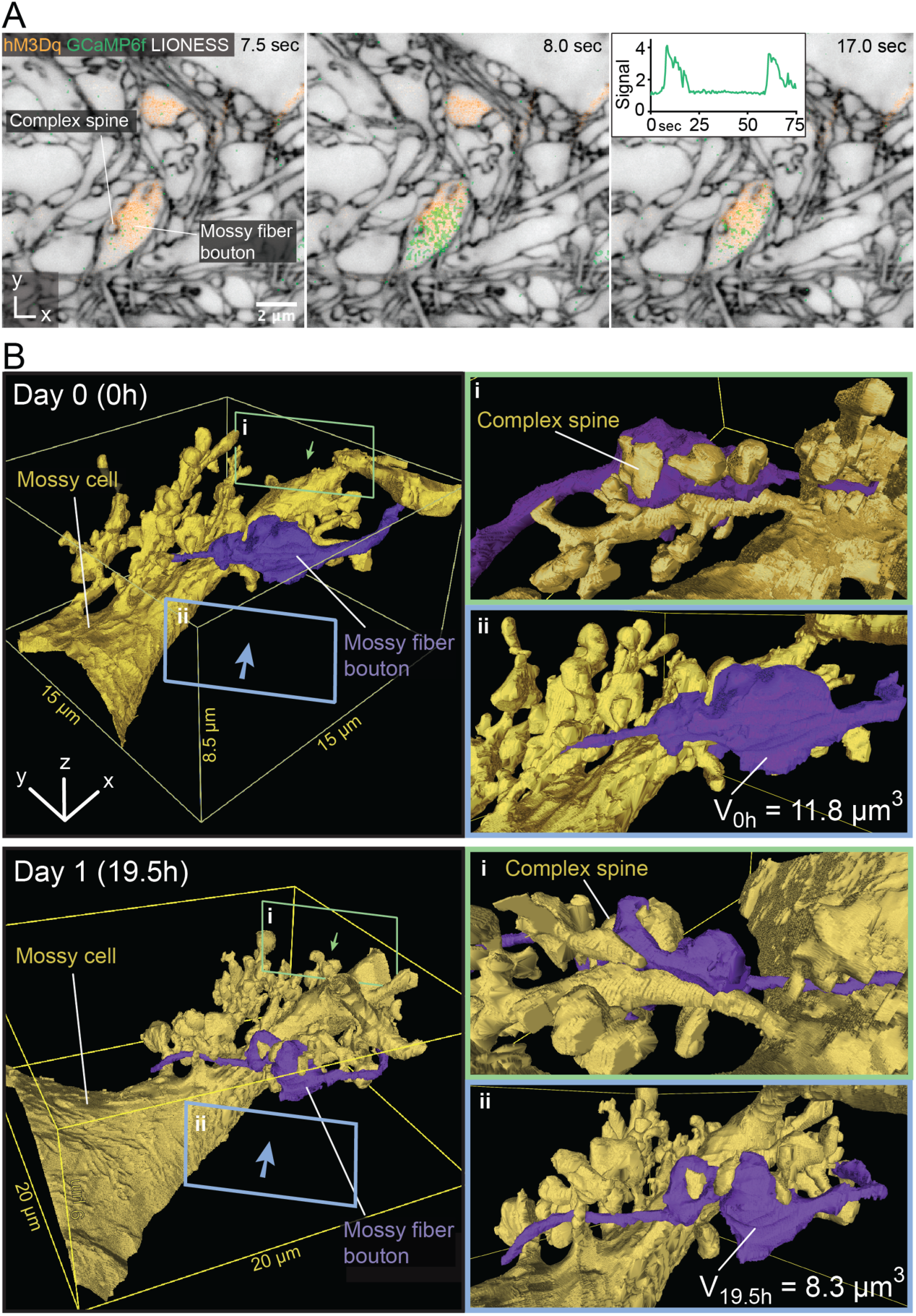
3D-morphodynamics and chemogenetically induced Ca^2+^-activity in hippocampal mossy fiber-hilar mossy cell synapses. (**A**) Single plane of a LIONESS volume in the hilus of the dentate gyrus in an organotypic hippocampal slice culture where a subset of mossy fiber boutons expressed both the excitatory DREADD hM3Dq (together with cytosolic dTomato, orange) and the calcium indicator GCaMP6f (green). LIONESS and dTomato images are identical replicates placing the overlaid time-varying Ca^2+^-signals after stimulation with the DREADD ligand CNO into structural context, with 3 exemplary points from a time series. Insert: GCaMP signal (averaged pixel value normalized to first frame) as a function of time. LIONESS image is a maximum intensity projection spanning 150 nm. Scale bar: 2 µm. (**B**) 3D-reconstructions of a hM3Dq-expressing mossy fiber bouton (purple) and its postsynaptic partner, a hilar mossy cell (gold) with complex spines at two timepoints (top: day 0 (0h), bottom: day 1 (19.5h)). V_0h_ and V_19.5h_ are bouton volumes at the respective time points. Green (i) and blue (ii) frames indicate the viewing angles from opposite directions for the magnified views on the right. The structures designated by the lettering in both panels refer to the same bouton and complex spine.

### Electrophysiology

We reasoned that with LIONESS, light microscopy may not only be used for visual guidance of electrophysiology experiments, but to correlate electrical properties of single and synaptically connected pairs of neurons with the underlying neuronal architecture in the living state. We performed whole-cell patch-clamp recordings of two neighboring pyramidal neurons in the hippocampal CA1 region, taking advantage of the fact that these cells often form monosynaptic connections in organotypic culture^42^. Cells with confirmed synaptic connectivity were imaged post-recording (**Suppl**. **Fig. 17)**, first in diffraction-limited mode with fluorophores intracellularly applied during recording, and then with LIONESS. Zooming in on a region of putative contact, diffraction-limited readout indicated that this was the site of electrophysiologically confirmed communication. Only comprehensive 3D-super-resolved delineation in the LIONESS volume revealed the deception by disclosing an intervening, unlabeled neuronal process missed in diffraction-limited mode. This corroborated that LIONESS was suitable for a multimodal approach to simultaneously retrieve and correlate structural with functional aspects of tissue architecture in living specimens, and more powerful in doing so than diffraction-limited imaging.

### Bridging scales

Analysis of tissue architecture was not limited to single LIONESS volumes. For extending analysis volumes and embedding them into meso-scale context, we followed two straight-forward approaches. Firstly, recording multiple partially overlapping subvolumes in the living tissue allowed automatically 3D-registering them with sufficient accuracy, such that segments from automated reconstruction smoothly extended over the borders. For example, we reconstructed a 70 μm-long stretch of mostly parallel axon fibers in acutely prepared mouse alveus from four image volumes – thus capturing ∼3 mm of cumulative axon length in this continuous region (**Suppl. Fig. 18**). Secondly, we guided selection of LIONESS volumes by recording larger volumes with diffraction-limited resolution, which provided further context to reconstructed regions. As one example, imaging a 650,000 µm^3^ volume in the DG crest of the hippocampus gave positional context and allowed identification of larger objects like cell somata and major dendritic branches, whereas LIONESS reconstruction revealed how DG granule cell and other cellular processes were embedded in the invaginations of a glial cell (**Suppl. Fig. 19, Suppl. Video 9, 10**). Imaging across spatial scales thus yields information on cell position and identity to extend the interpretation of connectivity analysis made possible by LIONESS.

## Discussion

Here we demonstrate saturated reconstruction of living mammalian brain tissue. This allowed tracking the time evolution of its structure, direct pairing with information on molecule location, and with simultaneous manipulation and readout of activity. Together, these elements constitute a fundamentally new quality of information for the study of brain structure and function, overcoming limitations of both static EM representations and reconstruction of positively labelled, incomplete subsets of tissue constituents in light microscopy.

To achieve this, we developed a technology for imaging and segmenting living brain tissue, integrating optimization of optical nanoscopy for tissue imaging with a two-step deep-learning strategy. It may come as a surprise that a moderate ∼130 nm isotropic 3D-resolution, chosen to limit burden on the living specimen, was sufficient for brain tissue reconstruction. This contrasts with ∼4x better resolution in the worst-resolved direction in typical EM reconstructions^1,2^, relying on physical sectioning at ∼30 nm steps. Four factors aided segmentation: 1) Extracellular labelling^14,17^ selectively highlighted the space separating cellular structures. The presence of a separating fluorophore layer was detectable also at resolution worse than the thickness of this layer. ECS labelling eclipsed complexity from intracellular organelles in the cell segmentation channel, whereas specific labelling revealed intracellular structures and molecules in spectrally distinct detection channels. 2) Also cellular structures smaller than the effective point-spread-function of the microscope led to modulation of fluorescence counts by volume exclusion of extracellular label, aiding in their detection. 3) Live imaging lacks alterations of tissue structure from chemical fixation, including shrinkage^43^ of the space filled with label in between cells. 4) Optical sectioning yields inherently aligned volumetric stacks, avoiding potential technical challenges with alignment after physical sectioning.

Our parameter search for image acquisition, processing, and segmentation was in no way exhaustive, opening possibilities for future improvements. For example, beyond brute-force further extension of the training bases for both deep-learning networks, the first network may be trained on paired low/high-exposure volumes where the high-exposure ground truth also features increased resolution. LIONESS will likely benefit from specifically engineered fluorophores for high performance extracellular labelling, which we have only started to explore by attaching hydrophilicity-enhancing sulfonate groups.

The capacity of LIONESS to bridge to the wider tissue context will benefit biological studies. Our chosen LIONESS volumes of up to ∼4,500 µm^3^ reflect best optical performance within a few tens of µm depth and in the central region of the objective’s field of view. We increased these laterally via volumetric tiling whereas adaptive optics^44^ may be used to enhance axial range. Rather than maximizing imaging volumes, one may also bridge scales with complementary methods, such as viral tracing. While we used rabies virus for live labelling of synapses, this approach may be extended to retrogradely trace^38,45^ the cellular origins of synaptic input identified in LIONESS. Application of LIONESS in living animals harbors engineering challenges due to movement^46^ that may be addressed by developing motion-compensating algorithms. We envision that correlating live information from LIONESS with light microscopy measurements of the same specimen after fixation will be useful, as this opens up further possibilities for molecular characterization and large-scale super-resolution tissue imaging^47^. Such all-optical correlative measurements will be considerably less complex than correlative approaches with EM. As expected, optimum sample and imaging conditions were required for full reconstruction. However, STED imaging was performed on a commercial microscope with minor adaptations, which underscores adoptability of this technology. The simplified workflow and lower demands in equipment or personnel of LIONESS compared to serial-sectioning or block-face imaging EM should enable analysis of multiple specimens as a function of genotype, developmental stage, disease, or specific intervention.

More fundamentally though, LIONESS is unique in that it allows repeated reconstruction of brain tissue over time, capturing single-synapse to network-level structural plasticity together with location of specific molecules and neuronal activity directly in the living state. For determining structure-function relationships, this sets it apart also from functional EM studies of brain tissue, including in correlative Ca^2+^ imaging^3,4^ and “flash and freeze” EM^48^. Taking advantage of extracellular rather than sparse cellular labelling, LIONESS resolves structures with their interacting partners. This provides resolution of both neuronal input and output, as well as tissue microenvironment, and facilitates observation of how activity and plasticity of one component translates to surrounding structures. Likewise, it offers new opportunities for studying synaptic heterogeneity, which underlies much of the complexity of the brain. Our results demonstrate that LIONESS is a powerful tool that can provide new biological insights into the function of neuronal circuits. As LIONESS can be applied to live tissue, it is possible to monitor synaptic connectivity over time, permitting real-time analysis of the dynamics of the connectome. Furthermore, since LIONESS can be applied in conjunction with molecular labels (such as PSD95), structural and functional connectivity can be analyzed in parallel. Thus, LIONESS may contribute to the identification of structural and functional components of synaptic engrams in neuronal circuits. Finally, because LIONESS is fully compatible with Ca^2+^-imaging, it allows to correlate synaptic plasticity with the pre- and postsynaptic activity history of the synapse, allowing future users to define the precise induction rules of synaptic plasticity and engram formation.

LIONESS has the capability to analyze the structure of living brain tissue in a comprehensive, and thus unbiased way. This provides rich opportunities for unexpected discoveries that may ultimately challenge the way we think about the extent and significance of plasticity in the central nervous system. We extracted exemplary quantitative data on neuronal connectivity and structure to showcase that LIONESS provides quantitative, dynamical biological information. We thus expect it to facilitate addressing biological questions related to stability and rewiring of network structure^49^, the dynamics of neuron-glia interactions and ECS^19,20^, and structure-function relationships^48^. LIONESS opens up new avenues for decoding complex, dynamic tissue architecture in living mammalian brain and other organs.

## Author contributions

P.V. designed and performed experiments, analysed, proofread, visualized and interpreted data, and prepared figures. E.M. set up and performed automated segmentation, data analysis, and visualization. J.M.M. supported experiments. D.W. and Z.L. supported automated segmentation. J.F.W. performed patch-clamp experiments and segmentation for validation. J.T. performed visualization, advised by J.Be.. Y.B.-S. provided viral constructs. C.S. supported image analysis. W.J. supported setting up imaging and troubleshooting. A.C. performed manual segmentations. J.Br. synthesized SulfoAtto 643. S.G.N.G. provided PSD95-HaloTag mouse. P.J. supervised patch clamp experiments and virus generation. G.N. advised on and provided human cerebral organoids. H.P. advised on automated segmentation. B.B. supervised computer vision. J.G.D. conceived and supervised the study, designed experiments, and interpreted data. J.G.D. wrote the paper together with P.V. with critical input from all authors.

## Acknowledgements

We thank J. Vorlaufer, N. Agudelo, A. Wartak for microscope maintenance and troubleshooting, C. Kreuzinger and A. Freeman for technical assistance, and J. Lyudchik and M. Šuplata for computational support and hardware control and Márcia Cunha dos Santos for initial exploration of software. We thank Paul Henderson for advice on deep-learning training and Michael Sixt, Scott Boyd, and Tamara Weiss for discussions and critical reading of the manuscript. Luke Lavis (Janelia Research Campus) generously provided JF585-HaloTag ligand. We acknowledge expert support by IST Austria’s scientific computing, bioimaging, preclinical, and life science facilities, and by the Miba machine shop. This work was supported with funding by the Austrian Science Fund (FWF) (I3600-B27 and DK W1232) to J.G.D. and (Z 312-B27, Wittgenstein award) to P.J., by the Gesellschaft für Forschungsförderung NÖ (NFB) to J.G.D. (LSC18-022), an institutional (ISTA) Interdisciplinary Project grant to J.G.D. and B.B., the European Research Council (ERC) under the European Union’s Horizon 2020 research and innovation programme to B.B. (715767 – MATERIALIZABLE), to G.N. (715508 – REVERSEAUTISM), to S.G. (695568 – SYNNOVATE), and to P.J. (692692 – GIANTSYN), the Simons Foundation Autism Research Initiative to S.G. (529085), and the Wellcome Trust to S.G. (Technology Development Grant 202932). It was further supported by a MSCA Individual Fellowship under the EU Horizon 2020 program to J.F.W. (101026635) and a postdoctoral fellowship from the Human Frontier Science Program (HFSP) to W.J. (LT000557/2018). This work was also partially supported by NSF grants IIS-1835231 and NCS-FO-2124179 to H.P..

## Methods

### Animals

Animals were housed in groups of 3–4 animals per cage and kept on a 12 h light/dark cycle (lights on at 7:00 am), with food and water available ad libitum. If not stated otherwise, we used wild-type C57BL/6J mice. All transgenic lines (see Table 1) used in this study have been previously characterized. For experimental use, we crossed Ai95 (GCaMP6f)^40^ and Prox1-cre. For experiments with PSD95-HaloTag mice^36,37^, both homozygous and heterozygous animals were used. For all experiments, male and female mice were used interchangeably. Experiments and procedures were performed in strict accordance with institutional, national, and European guidelines for animal experimentation.

**Table 1.**

| Transgenic line | Full name | Source (deposited by) | Cat# |
| --- | --- | --- | --- |
| Ai95 | B6;129S-Gt(ROSA)26Sortm95.1(CAG-GCaMP6f)Hze/J | Jackson labs (H. Zeng) | 024105 |
| Prox1-cre | Tg(Prox1-cre)SJ32Gsat/Mmucd | MMRRC (N. Heintz) | 036644-UCD |
| PSD95-HaloTag | PSD95-HaloTag | Seth G. N. Grant, Edinburgh | - |
| Thy1-EGFP<sup>50</sup> | STOCK Tg(Thy1-EGFP)MJrs/J | Jackson labs (J. Sanes) | 007788 |

### Organotypic hippocampal slice cultures

Hippocampal slices were obtained from 5-7 days old mice of either sex and cultured on cell culture inserts with porous membranes. Mouse pups were decapitated and the hippocampus was isolated while the brain was submerged in ice cold sterile filtered HBSS without Ca^2+^ and Mg^2+^ (Gibco, #14175-053) supplemented with 10 mM glucose, using a stereo microscope. Hippocampi were cut into 350 µm thick slices and placed on round porous membranes with 4 mm diameter (PTFE membrane, Merck, #FHLC01300), which have been placed on cell culture inserts with a porous membrane (Millicell, #PICM0RG50) for interface culture. The inserts with the slices were placed in dishes (Greiner, #627161) with 1 ml of culture media. We adapted the media recipe during the course of experiments, as quality of cultures deteriorated with the same nominal composition. We found 78.5% Minimum Essential Medium (MEM, Gibco, #11095-080), 15% heat-inactivated horse serum (Gibco, #26050070), 2% B27 supplement (Gibco, #0080085SA), 2.5% 1 M HEPES (Sigma, #M3375-100G), 1.5% 0.2 M GlutaMax supplement (Gibco, #35050-061), 0.5% 0.05 M ascorbic acid (Sigma, #A5960-25G), with additional 1 mM CaCl_2_ and 1 mM MgSO_4_ to produce satisfactory results, and incubated at 37 °C and 5% CO_2_. The medium was changed the day after preparation and then every 3-4 days.

### ECS labelling

For ECS labelling, artificial cerebrospinal fluid (ACSF) was prepared from a 10x stock solution with MgCl_2_ and CaCl_2_ added freshly before carbogen bubbling, whereas ascorbic acid and Trolox were added after bubbling. Finally, ACSF consisted of 125 mM NaCl, 2 mM CaCl_2_, 1.3 mM MgCl_2_, 4.8 mM KCl, 26 mM NaHCO_3_, 1.25 mM NaH_2_PO_4_, 7.5 mM HEPES (Gibco, #15630056), 20 mM D-glucose (Sigma, #G8270-1kg), 1 mM Trolox (Sigma, #238813), 1 mM ascorbic acid (Sigma, #A5960-25G); pH 7.4. Thereafter, fluorescent dye (Atto 643 (Atto-Tec GmbH, #AD 643-25), SulfoAtto 643, or Abberior STAR 635P (Abberior, #ST635P)) was added from 5 mM stocks (dissolved in ACSF) to a final concentration of 150 µM. A 2 µl droplet of the dye-containing imaging solution was put on a #1.5H coverslip (Bartelt, #6.259 995) that had been placed in an imaging chamber (RC-41, Warner Instruments). Using fine forceps, brain slices with the membrane attached were then carefully put onto the droplet, such that the slice was oriented towards the coverslip. A slice anchor gently kept the sample in place. Immediately afterwards, further imaging solution at room temperature (RT) was added. The imaging chamber was then placed onto the stage adapter of the STED microscope (see below). The data in the manuscript were acquired using Atto 643, except for **Fig. 5** and **Suppl. Fig. 11** (left panel) where SulfoAtto 643 was used.

### Acute preparation of whole hippocampus and labelling

Hippocampi were extracted from 5-7 days old mice of either sex. Mouse pups were decapitated and the hippocampus isolated while the brain was submerged in ice cold sterile filtered HBSS without Ca^2+^ and Mg^2+^ (Gibco, #14175-053) supplemented with 10 mM glucose, using a stereo microscope. The whole hippocampus was then submerged in freshly carbogenized ACSF with 150 µM Atto 643 dye and incubated for 10 min at RT with gentle agitation. Afterwards, entire hippocampi were placed on a #1.5H coverslip that had been placed in an imaging chamber (RC-41, Warner Instruments) with the alveus region facing the coverslip. A slice anchor gently kept the sample in place when freshly carbogenized ACSF with 150 µM Atto 643 dye was added for imaging. The imaging chamber was then placed onto the stage adapter of the STED microscope.

### Generation of cerebral organoids

Human embryonic stem cells were dissociated to single cells using Accutase (Gibco). A total of 2500 cells was transferred to each well of an ultra-low-binding-96-well plate (Corning) in mTeSR1 media supplemented with 50 µM Y-27632 (Stemcell Technologies). Cells were allowed to aggregate to EBs and fed every second day. At day 3 supplements were removed and from day 6 the generation of cerebral organoids was performed according to Lancaster and Knoblich^34^. Briefly, EBs were transferred to neural induction medium (NIM) in low-adhesion 24-well plates (Corning), and fed every second day for 5 days until formation of neuroepithelial tissue (day 0 of cerebral organoid formation). Neuroepithelial tissue-displaying organoids were embedded in Matrigel droplets (Corning, #356234) and grown in cerebral organoid medium (COM) supplemented with B27 without vitamin A (Gibco) and fed every other day. After 4 days tissues were transferred to COM supplemented with B27 containing vitamin A and placed on a horizontal shaker at 70-100rpm. Cerebral organoids were fed twice a week.

### LIONESS imaging

STED microscopy was performed at room temperature on an inverted Expert Line STED microscope (Abberior Instruments) with pulsed excitation and STED lasers. A 640 nm laser was used for excitation and a 775 nm laser for stimulated emission. A silicone oil immersion objective with 1.35 NA and a correction collar (Olympus, UPLSAPS 100XS) was used for image acquisition. The fluorescence signal was collected in a confocal arrangement with a pinhole size of 0.6 or 0.8 airy units. For detection a 685/70 nm bandpass filter (Chroma, #F49-686) was used and a 50:50 beam splitter (Thorlabs, #BSW29R) distributed the signal onto two photon counting avalanche photodiodes, allowing for stronger excitation without saturating detectors. Both detection channels were added up using Fiji^51^ Version: 2.3.0/1.53f (Fiji/process/calculator plus/add), photon counts inverted, and data saved in 16-bit TIFF format. The pulse repetition rate was 40 MHz and fluorescence detection was time-gated. LIONESS volumes were acquired with 10 µs pixel dwell time, 2.9 µW (640 nm) excitation laser power and 90 mW STED laser power. A spatial light modulator (SLM) imprinted incoherently overlapped phase patterns for predominantly axial resolution increase (π-top-hat phase modulation, z-STED), and for predominantly improved fluorescence quenching outside the central minimum (4π-helical phase modulation, 4π(*xy*)-STED) onto the STED beam. The SLM was also used to perform alignment directly in the sample, ensuring that the intensity minima of the two STED patterns spatially coincided and to optionally adjust low-order Zernicke polynomials for empirical aberration correction. Power ratio of *z*-STED/xy(4)-STED/ was 80/20. Voxel size was 50 x 50 x 50 nm^3^ for all images. Acquisition scan mode was typically *xzy*, with the *y*-direction being the slowest scan axis, using galvanometric mirrors for lateral (*xy*)-scanning and a sample piezo stage (Physik Instrumente (PI) GmbH & Co. KG, #P-736.ZRO) for axial (*z*)-scanning. Image acquisition and microscope control were performed with Imspector software version 14.0.3052.

For samples with additional positive labels (HaloTag ligand JF585, Synaptophysin-EGFP, Thy-1-EGFP, GCaMP6f), additional color channels with diffraction-limited resolution using a 488 nm or 560 nm laser with 10 µs dwell time and 1.1- 3.9 µW (488 nm) and 2- 2.6 µW (560 nm) excitation power were used for recordings. These signals were collected using a photon counting avalanche photodiode with a 525/50 nm (Semrock, #F37-516) and 605/50 nm (Chroma, #F49-605) bandpass filters for EGFP and JF585 detection, respectively. The 488 nm and 640 nm excitations were done simultaneously, for 560 nm excitation a second line step was used to avoid spectral bleed-through into the far-red channel. Voxel size was again 50 x 50 x 50 nm^3^ for all images with *xzy*-scan mode. The power values refer to the power at the sample, measured with a slide powermeter head (Thorlabs, S170C).

### Repeated volumetric live imaging

For evaluation of tissue photo-burden with LIONESS vs. conventional high-exposure STED (**Suppl. Fig. 4**) a 70 x 70 µm confocal overview scan was performed in a region of neuropil in the CA1 region of an organotypic hippocampal slice. Next, the central 5 x 5 x 2.5 µm^3^ volume was exposed to STED in 20 consecutive volumetric scans in *xyz*-scan mode with 70 µs voxel dwell time for long-exposure STED and 10 µs for LIONESS datasets. Excitation and STED power were identical and corresponded to the parameters used in LIONESS imaging, with 90 mW STED power at 80/20 distribution between phase patterns. 10 min after the last volume was acquired, a second 70 x 70 µm confocal overview scan was done of the same region and plane as in the initial measurement.

For long-term repeated imaging of hippocampal neuropil (**Suppl. Fig. 15**), the sample was mounted and placed on the microscope as described in the section on LIONESS imaging. For the first 4 acquisitions within 1h, the sample was kept in place, with the imaging media (carbogenized ACSF with 150 µM Atto 643) exchanged after 30 min. After that, the sample was placed back onto cell culture inserts and into the tissue culture incubator at 37 °C and 5% CO_2_ until the next imaging session one day later. The same procedure was repeated for the last imaging time point after 3 days.

For long-term repeated imaging of chemogenetically activated mossy fiber boutons (**Fig. 5**), the sample was placed back after the first imaging session onto cell culture inserts and incubated at 37°C and 5% CO_2_. Media was changed after 45 min to wash out residual CNO, and the sample was placed into the tissue culture incubator until the second imaging session on the next day.

### PSD95-HaloTag labelling

Organotypic hippocampal brain slices of PSD95-HaloTag^36,37,52^ mice were live labelled using Janelia Fluor (JF)585-HaloTag ligand (Janelia Research Campus). The fluorescent ligand was dissolved in anhydrous DMSO to a stock concentration of 500 µM, aliquoted and stored at -20°C. Before imaging, the fluorescent ligand was added to the culture medium to a final concentration of 500 nM (1:1000) and incubated for at least 45 min at 37 °C.

### Viral vector assembly and synaptophysin labelling

Preparation of AAV and RVdG_envA_-CVS-N2c vectors has previously been described^38,53^. Briefly, AAV2-CaMKIIa-TVA-2A-N2cG (Addgene #172363) vectors were pseudotyped with the AAVdj capsid protein by co-transfection of HEK293T cells. Three days later, the cells were harvested and lysed, and the viral stock was purified using heparin-agarose affinity binding. RVdG_envA_-CVS-N2c-nl. EGFP-SypEGFP (Addgene #172380) were rescued using HEK-GT cells and then amplified and pseudotyped using BHK-eT cells. Viral vectors were purified and concentrated from the supernatant using ultracentrifugation and resuspended in PBS.

For live labelling of synaptic vesicles, first AAV-CaMKIIa-TVA-2A-N2cG was added to organotypic hippocampal slice cultures at 7-10 days in vitro (DIV) for dual expression of the TVA avian receptor and the rabies N2c glycoprotein (N2cG). 14 days later, envA-pseudotyped, G-deleted CVS-N2c rabies viral particles were added for expression of a synaptophysin-EGFP fusion protein and additional EGFP expression in the cell nucleus (RVdG(envA)-CVS-N2c-nlGFP-sypGFP). 4-5 days after addition of the rabies vectors, EGFP expression was strong enough for imaging.

### Myelin labelling

Live labelling of myelin was performed using FluoroMyelin^TM^ Green (ThermoFisher Scientific, #F34651). The dye was diluted 1:300 in culture media for organotypic hippocampal slices and incubated with the sample at 37 °C for at least 30 min before imaging.

### Calcium imaging

Cultured organotypic hippocampal slices of Prox1-cre/Ai95 (GCaMP6f)^40^ mice shown in **Suppl. Fig. 16** were ECS labelled for LIONESS imaging as described above. To reduce level of inhibition, 10 µM GABAA receptor antagonist gabazine were added to the imaging media at the start of the imaging session. A region of interest was first repeatedly imaged via confocal scanning (488 nm excitation, 1.1 µW) of an individual plane with 50 x 50 nm2 pixel size and 5 µs pixel dwell time (frame rate ∼1.25 Hz) to detect GCaMP signals. After recording, the enclosing volume was scanned in LIONESS mode. The GCaMP recording was overlaid with a corresponding plane of the volumetric LIONESS acquisition in **Suppl. Fig. 16C**. The same volume was imaged a second time 10 min after the first acquisition. The sample was kept in place in between the two recordings.

### Chemogenetic activation with calcium imaging

Chemically targeted activation with simultaneous calcium imaging of neurons was done using AAVs containing a Cre-dependent DREADD^54^ construct (AAV-DIO-CAG-hM3Dq-2A-dTomato; plasmid available from the authors upon request) added to organotypic hippocampal slice cultures of Prox1-cre/Ai95 (GCaMP6f) mice at DIV 4-6. Each transduced cell expressed both cytoplasmic dTomato and the excitatory designer receptor hM3Dq. Concentrated viral stock (7×10^11^ GC/ml) was first diluted 1:10 in culture medium, and subsequently 5 µl were carefully placed on top of each slice. Weak fluorescence was already detectable ∼3 days after transfection and live-imaging was performed from day 9 onwards after viral transduction. To activate the designer receptor, Clozapine-N-oxide (CNO) was added (3 µM final concentration) to the imaging medium (fluorophore containing ACSF). The GCaMP signal was recorded via confocal scanning (488 nm excitation, 3.9 µW) of an individual plane using a pixel size of 100 x 100 nm^2^ and dwell time of 20 µs, which resulted in a frame rate of ∼ 2 Hz. The GCaMP recording together with the dTomato signal were overlaid with a corresponding plane of the LIONESS acquisition for representation. For the inset in **Fig. 5A**, a square region of interest around the CNO activated mossy fiber bouton was defined, GCaMP signal averaged over this region, and normalized to the value in the first frame.

### Electrophysiology

Organotypic slice cultures were submerged in artificial cerebrospinal fluid (ACSF) containing 125 mM NaCl, 25 mM NaHCO_3_, 25 mM D-glucose, 2.5 mM KCl, 1.25 mM NaH_2_PO_4_, 2 mM CaCl_2_, and 1 mM MgCl_2_, with pH maintained at 7.3, equilibrated with a 95% O_2_/5% CO_2_ gas mixture at ∼22 °C (room temperature). Glass micropipettes were pulled from thick-walled borosilicate glass (2 mm O.D., 1 mm I.D.) and filled with intracellular solution containing 135 mM K-gluconate (Sigma, #G4500), 20 mM KCl, 0.1 mM EGTA (Sigma, #E0396), 2 mM MgCl_2_, 4 mM Na_2_ATP (Sigma, #A3377), 0.3 mM GTP (Sigma, #G8877), 10 mM HEPES (Gibco, #15630056), with the addition of 20 µM AlexaFluor488 hydrazide (Invitrogen, #A10436) and 0.2 % (w/v) biocytin (Invitrogen, #B1592) as required. Pipettes were positioned using two LN mini 25 micromanipulators (Luigs and Neumann) under visual control on a modified Olympus BX51 microscope equipped with a 60x water-immersion objective (LUMPlan FI/IR, NA = 0.90, Olympus, 2.05 mm working distance). Two neurons were simultaneously recorded in the whole-cell patch-clamp configuration, with signals acquired on a Multiclamp 700B amplifier (Molecular Devices), low pass filtered at 6 kHz and digitized at 20 kHz with a Cambridge Electronic Design 1401 mkII AD/DA converter. Signals were acquired using Signal 6.0 software (CED). Action potential phenotypes were recorded on sequential current pulse injections (−100 to +400 pA) in the current clamp configuration. Neurons were identified based on morphological and action potential phenotypes. In current clamp recordings, pipette capacitance was 70 % compensated.

Synaptic connectivity was assessed by sequential current injection into either recorded cell in the current-clamp configuration, while recording EPSCs (Excitatory Postsynaptic Currents) from the other in the voltage-clamp configuration. Presynaptic action potentials were elicited by five 1-2 nA current injection pulses for 2-3 ms at 20 Hz. Putative monosynaptic connections were identified by EPSC generation (peak current > 2.5 times the standard deviation of baseline noise) in the postsynaptic cell with short latency (< 4 ms) from the presynaptic action potential peak. Recordings were analysed using Stimfit^55^ and MATLAB-based scripts.

After recording, neurons were resealed by forming an outside-out patch on pipette retraction, before immersion in solutions for live imaging.

### SulfoAtto 643 synthesis and characterization

In a 5 mL round bottom flask equipped with a magnetic stir bar, Atto 643 NHS-ester (ATTO-TEC: #AD 643-35; 5.0 mg, 5.23 µmol, 1.0 equiv.) was dissolved in a mixture of 700 µL *N*,*N*-dimethylformamide (Fisher Scientific: #D/3846/17) and 300 µL dH_2_O. *N*,*N*-Di*iso*propylethylamine (Carl Roth: #2474.1) (6.9 mg, 53.8 µmol, 9.3 µL, 10 equiv.) and taurine (Carl Roth: #4721.1) (3.4 mg, 26.8 µmol, 5.1 equiv.) were added successively and the reaction mixture was allowed to incubate under stirring for 60 min before it was quenched by the addition of glacial acetic acid (Carl Roth: #6755.1) (10 µL). Semi-preparative reverse phase-high pressure liquid chromatography was performed on an Agilent 1260 Infinity II LC System equipped with a Reprospher 100 C18 column (5 µm: 250 x 10 mm at 4 mL/min flow rate). Eluents A (0.1% trifluoroacetic acid (TCI: #T0431) in dH_2_O) and B (0.1% trifluoroacetic acid in acetonitrile (Honeywell: #34851-2.5L)) were used. The gradient was from 10% B for 5 min → gradient to 90% B over 35 min → 90% B for 5 min with 4.0 mL/min flow. Peak detection and collection were performed at λ = 650 nm and provided 4.5 mg (4.7 µmol) of the desired product as a blue powder after lyophilization with 91% yield. Characterization was performed using high pressure liquid chromatography mass spectrometry (**Suppl. Fig. 20, left**) on an Agilent 1260 Infinity II LC System equipped with Agilent SB-C18 column (1.8 µm, 2.1 × 50 mm). Buffer A: 0.1% formic acid (Fisher Scientific: A117-50) in dH_2_O Buffer B: 0.1% formic acid in acetonitrile. The gradient was from 10% B for 0.5 min → gradient to 95% B over 5 min → 95% B for 0.5 min → gradient to 99% B over 1 min with 0.8 mL/min flow. Retention time *t_R_* = 3.03 min. Low resolution mass spectrometry: calculated: 943 Da, found: 943 Da. Excitation and emission spectra were recorded on a TECAN INFINITE M PLEX plate reader (λ_Ex_ = 580±10 nm; λ_Em_ = 620–800±20 nm; 10 flashes; 40 µs integration time; λ_Ex_ = 300–660±10 nm; λ_Em_ = 700±20 nm; 10 flashes; 40 µs integration time) with 200 nM solutions of SulfoAtto 643 in PBS (Carl Roth: #9143.2) in Greiner black flat bottom 96 well plates (Carl Roth: #CEK8.1) (**Suppl. Fig. 20, right**).

### Restoration network training

Volumetric paired low-exposure, low-SNR training input data and high-exposure, high-SNR “ground truth” data were recorded in a voxel-exact mode by collecting low-SNR data during the first 10 µs voxel dwell time and additional photons during the remaining 60 µs dwell time. High-SNR ground truth for network training were thus generated by adding up counts from the total 70 µs dwell time in FIJI Version: 2.3.0/1.53f (Fiji/process/calculator plus/add). Other imaging parameters were as described in the section “LIONESS imaging” (2.9 µW (640 nm) excitation laser power, 90 mW STED laser power with power ratio of *z*-STED/*xy*(4)-STED of 80/20, voxel size 50 x 50 x 50 nm^3^). 76 volume pairs of 12.5 x 12.5 x 5 µm each were used for training. Volumes were taken from organotypic hippocampal and cerebellar slice cultures and the alveus region of acutely dissected hippocampi. Network training (Version: CSBDeep 0.6.1)^27^ parameters were as follows: 3D mode, 32 x 32 x 32 pixel patch size, 190 patches per volume, 150 steps per epoch, 150 epochs, batch size 32, and training data was loaded as 16-bit TIFF files. Software was installed from GitHub (https://github.com/CSBDeep/CSBDeep). A workstation with the following hardware components was used: Intel® Xeon® W “Skylake” W-2145, 3.60 GHz processor, 128 GB RAM, NVIDA GeForce RTX 2080Ti graphics card.

### Denoising

To denoise confocal images recorded simultaneously with the LIONESS data in **Fig. 4** and **Suppl. Fig. 6** and **14B**, Noise2void^56^ (Version 0.2.1) was applied to individual channels with the following parameters: noise2void 3D mode, patch size 32 x 32 x 32 pixels, each patch augmented with rotations and axis-mirroring, training steps per epoch 150, number of epochs 75 (SYP1-EGFP) or 100 (PSD95-HaloTag), batch size 16 (SYP1-EGFP) or 32 (PSD95-HaloTag). Software was installed from GitHub (https://github.com/juglab/n2v). A workstation with the following hardware components was used: Intel® Xeon® W “Skylake” W-2145, 3.60 GHz processor, 128 GB RAM, NVIDA GeForce RTX 2080Ti graphics card.

### Image analysis and processing

All used LUTs were linear except for **Fig. 2**, **Suppl. Figs. 2A-C, 5**, and **11**, where a color calibration bar is provided. Threshold adjustments for display purposes were applied linearly and to the whole image. Line profiles (**Fig. 1D** and **Suppl. Figs. 2C and 3**) were created using Fiji, line width was 2 pixels.

### Volume extension

For stitching of volumetric images, the Fiji 3D stitcher was used (Fiji/Plugins/deprecated/3D Stitching; linear blending, fusion alpha 2.0).

### Manual segmentation and proofreading

Planes for manual segmentation were first upscaled 5-fold without interpolation (plane depth was kept at original 50 nm spacing). Segmentation itself was done using VAST^57^ 1.3.0. and 1.4.0. Software was downloaded from https://lichtman.rc.fas.harvard.edu/vast/. For proofreading of automated segmentations data was visualized using Neuroglancer (https://github.com/google/neuroglancer) and corrected using VAST 1.4.0.

### Segmentation

We based our implementation of the automatic segmentation pipeline on the *pytorch_connectomics*^32,33^ framework. We used a U-Net architecture and trained the neural network to produce affinity maps which were then processed by a watershed algorithm to obtain the final segmentations.

During training, the U-Net required volume data and the corresponding manual ground-truth segmentation. First, in order to adapt the input datasets to the framework requirements and maximize its performance, we applied a pre-processing step converting the volume data to 8-bit format and stretching the intensity to cover the whole intensity range. Then, the pre-processed volumes together with the corresponding ground-truth segmentations were passed into the U-Net. Three key parameters during training were the sample size, the number of training iterations and the data augmentation. Given the anisotropic step size (5-fold upsampling in the *xz*- or *yz*-plane for manual segmentation) of the input volume we noticed that using a sample size of [128 x 128 x 64], with the lowest number corresponding to the non-upsampled axis, significantly improved the performance of the neural network over smaller sizes. We increased the number of training iterations from the default 100k to 500k which further helped reduce segmentation errors. We found this number of iterations to be a reasonable compromise between training time and inference performance. Finally, we enabled all available data-augmentation techniques.

During inference, we passed the pre-processed volume data into the U-Net and obtained the affinity map as output. At inference time we used the same sample size used during training, with appropriate padding if the input volume was small, and test-time augmentation via axis-mirroring. The values in the final affinity map corresponded to the mean of the values obtained for each augmented case. The output affinity map was processed using the watershed algorithm to produce the labelled automatic segmentation. Our pipeline combined two different watershed implementations. First, we applied the image-based watershed method^58^ (https://github.com/zudi-lin/zwatershed) on each slice to compute fragment masks. These were then passed to a volume-based implementation (https://github.com/zudi-lin/waterz), which was applied on the affinity map, producing the final segmentation. We used watershed thresholds in the range [0.2-0.4] to minimize oversegmentation but also avoid merges, which tend to be more tedious to fix during proof-reading. The resulting segmentations contained spurious segments, which we cleaned during a final post-processing step by removing those that consisted of too few voxels (fewer than 10) or slices (fewer that 2). This last step significantly facilitated later proof-reading. The resulting segmentations were then analyzed visually, using Neuroglancer, and quantitatively, using metrics such as segment size distribution and split ratio of ground truth segments with respect to automatic segmentations.

We trained the U-Net on a 8-GPU (NVIDIA 3090s) node, using 32 CPUs and 128 GB RAM during 500k iterations, which took 6 days. Inference time falls in the 10 to 40 minutes range, depending on the size of the input volume, and can be performed on a more modest compute node. In our case, we used a 2-GPU (NVIDIA 3090s) node using 8 CPUs. The post-inference watershed and segmentation cleaning operations were performed on the inference node and took 10 minutes to 20 minutes to complete.

### Visualization

3D visualizations were done either using VAST^57^ 1.4.0. (**Fig.1A, 2A** and **Suppl. Figs. 7, 8A, 11, 12, 19A**), Neuroglancer (**Fig. 3B** **, 4B, 5B, Suppl. Figs. 9, 10, 18, 19B**), Blender 2.93.4 (**Fig 1A, 3C, Suppl Fig. 8B**) or Neuromorph^59^ 2.8 (**Fig. 3E**). Blender-generated visualizations were produced based on 3D meshes extracted from segmentations using marching cubes (as implemented in Scikit-Image). These 3D meshes were first smoothed in Blender using a vertex-based smoothing operation that flattens angles of mesh vertices and finally the scene was rendered using Blender’s Cycles rendering engine. The schematics in the upper row of **Fig. 1A** were created with Biorender.com.

### Dendrite abstraction

For representing dendrite synaptic connectivity in **Fig. 3E**, we developed a visual spine analysis approach inspired by Barrio^60^, a software for visual neighborhood analysis of nanoscale neuronal structures. We computed surface meshes for all axons and dendrites based on the segmented neuronal structures. Next, we used Neuromorph^59^ to compute spine lengths by specifying the base and tip of each spine and plotted spine positions and relative spine length according to position on the dendrite. Spine lengths were computed between the base and tip of each spine, following the spine’s central axis (skeleton). We abstracted the complex 3D morphology and connectivity of a dendrite from 3D to 2D to reduce visual clutter, while preserving relative spine positions and spine lengths. To do so, we mapped a dendrite’s 3D skeleton structure to a simplified, but topologically correct, 2D representation. We preserved all relative distance relations within a dendrite (i.e., distances between spines), and encoded spine length at each spine location. Spine lengths were represented as bars, scaled relatively to the largest spine length of the dendrite.

### Statistics and reproducibility

In all images, representative data from single experiments are shown. LIONESS imaging of cerebral organoids as depicted in **Fig. 1A** and **Suppl. Fig. 8** was additionally repeated on similar specimens twice. Acute preparation of hippocampus and LIONESS imaging of the alveus region as shown in **Suppl. Figs. 10 and 18** was repeated 4 times, neuropil of wild type organotypic hippocampal slice cultures as in **Fig. 3** and **Suppl. Figs. 1, 2, 5, 12** and **19** was repeated at least 20 times. LIONESS imaging paired with PSD95-HalTag/SYP1-EGFP live labelling as in **Fig. 4** and **Suppl. Fig. 14** was repeated 4 times. Repeated imaging of the same region using LIONESS as in **Fig. 5** and **Suppl. Figs. 4b, 15 and 16** was performed at least 4 times. Repeated imaging of the same region with conventional, high photon load STED as shown in **Suppl. Fig. 4A** was repeated with performing *xy*-scanning only, showing the same negative effect. Comparison of single versus split detection (**Suppl. Fig. 3**) and 2π- and 4π-helical phase modulation (**Suppl. Fig. 2**) were repeated at least 3 times each, the comparison between single detection and 2π-helical phase modulation with split detection and 4π-helical phase modulation (**Fig. 1B, C**) was repeated 3 times. Fluoromyelin labelling as shown in **Suppl. Fig. 13** was performed 3 times.

---

**Suppl. Fig. 1.**
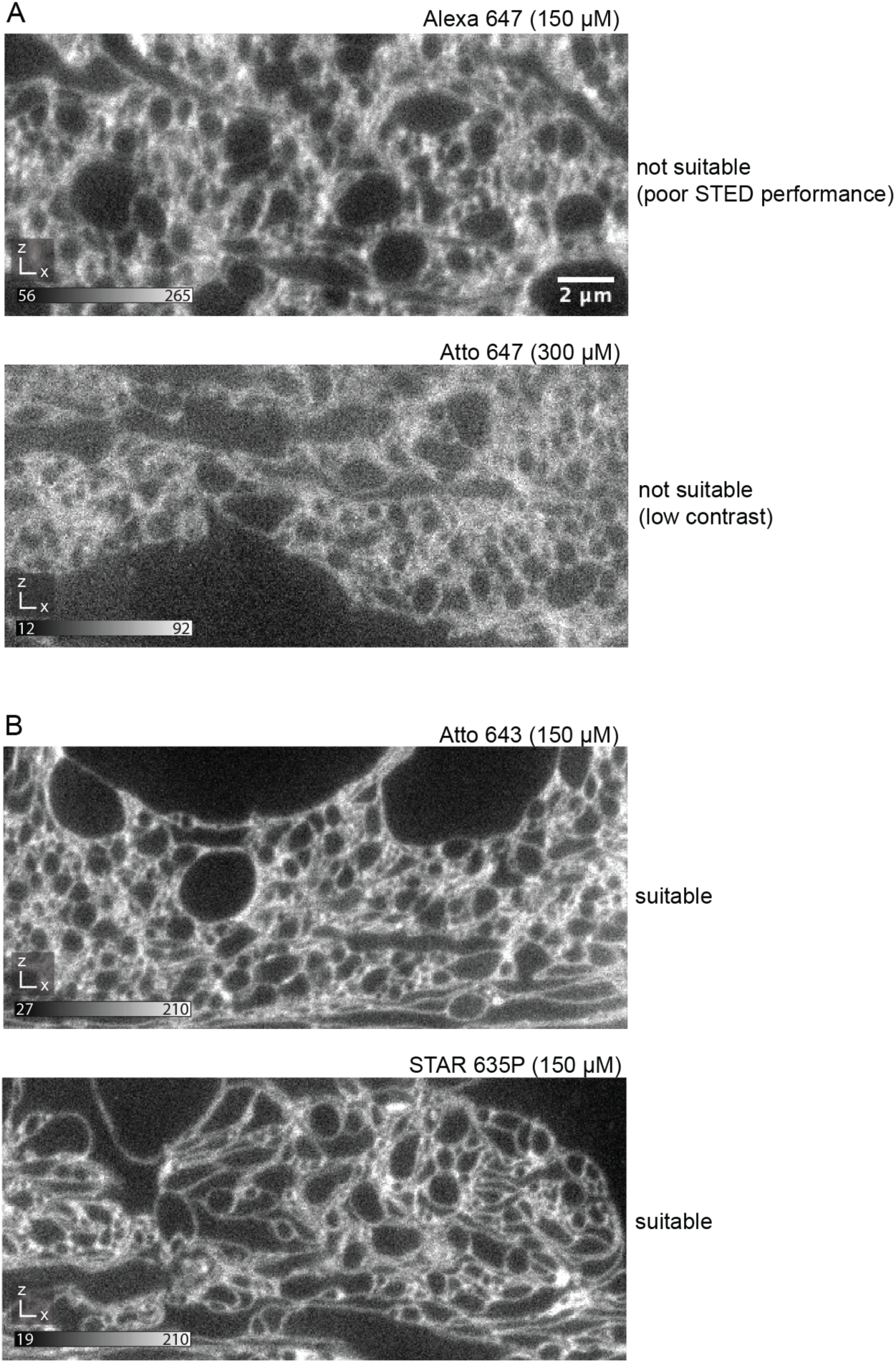
Fluorophore screening. (**A**) Two examples of fluorophores yielding insufficient delineation of fine cellular structures, due to suboptimal STED performance (top) or poor extra- vs. intracellular contrast (bottom). (**B**) Two examples of fluorophores with high STED performance and high extra- vs. intracellular contrast (Atto 643, Abberior STAR 635P), yielding adequate delineation of fine cellular structures. All images show raw *xz*-planes recorded with tissue-optimized STED patterns at near-isotropic resolution. Fluorophores applied at indicated concentrations to ECS in organotypic hippocampal slice cultures. The custom synthesized sulfonated variant of Atto 643 (SulfoAtto 643) was equally suited and used interchangeably with Atto 643. Scale bar: 2 µm, valid for all images. Numbers in greyscale bars refer to raw photon counts.

**Suppl. Fig. 2.**
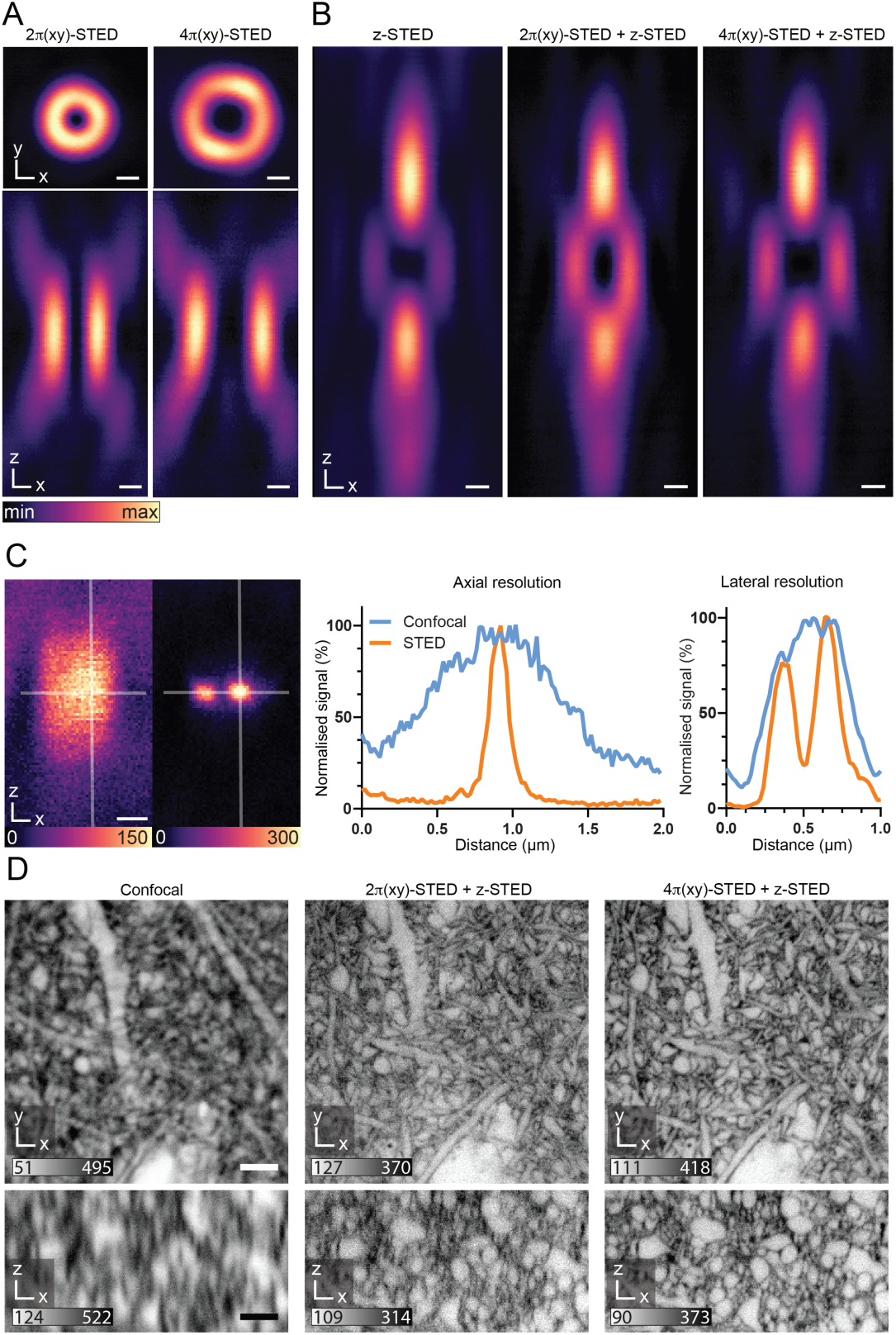
Tissue-optimized STED. (**A,B**) STED light intensity distributions in the focal region. (**A**) Lateral (top) and axial (bottom) sections for 2 - and 4 -helical phase modulation. Scale bars: 250 nm. (**B**) Axial sections of the -top-hat phase modulated *z*-STED pattern (left), an incoherent superposition of 2 (*xy*)- and *z*-STED patterns (middle), and of 4 (*xy*)- and *z*-STED patterns (right). Power distribution between the *z*- and *xy*-STED patterns in the superpositions was 80% vs. 20%. Scale bars: 250 nm. (**C**) Axial scan of 40 nm diameter fluorescent beads in confocal mode (left) and with STED employing combined 4 (*xy*)- and *z*-STED patterns (right). Scale bar: 250 nm. Profiles along the lines in lateral and axial directions as indicated in the images. (**D**) Extracellularly labelled neuropil in organotypic hippocampal slices. Orthogonal planes in *xy*- and *xz*-direction for diffraction-limited confocal (left), classical 2 -helical and -top-hat phase modulation (middle), and combination of 4 -helical plus π-top-hat modulation for near-isotropic resolution with improved quenching of excitation outside the central STED intensity minimum. The 4π-helical pattern also facilitated robust in-tissue co-alignment of intensity minima. Scale bar: 2 µm. Raw data with linear, inverted color scale. Numbers in greyscale bars refer to raw photon counts.

**Suppl. Fig. 3.**
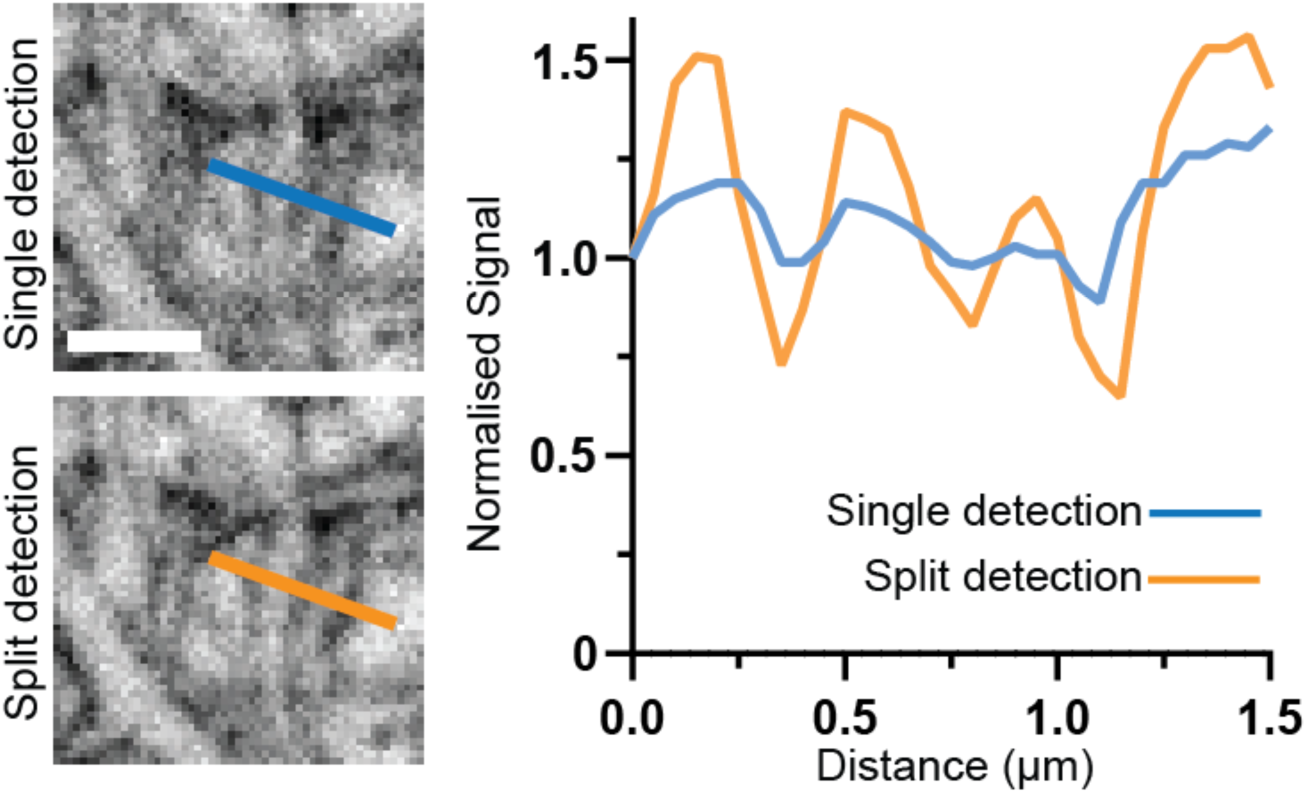
Detector dynamic range. Same region in organotypic hippocampal slice culture imaged with a single detector (top) or with a split detection path and two single-photon counting avalanche photodiodes as detectors (bottom). Line profiles over corresponding structures for single (blue) and split (orange) detection, normalized to first data point. STED power and pixel dwell times were identical, and STED patterns for near-isotropic resolution were used. Increased detector dynamic range allowed doubled excitation power within the linear detection regime, improving signal-to-noise ratio. Single imaging planes in organotypic hippocampal slice cultures. Scale bar: 1 µm.

**Suppl. Fig. 4.**
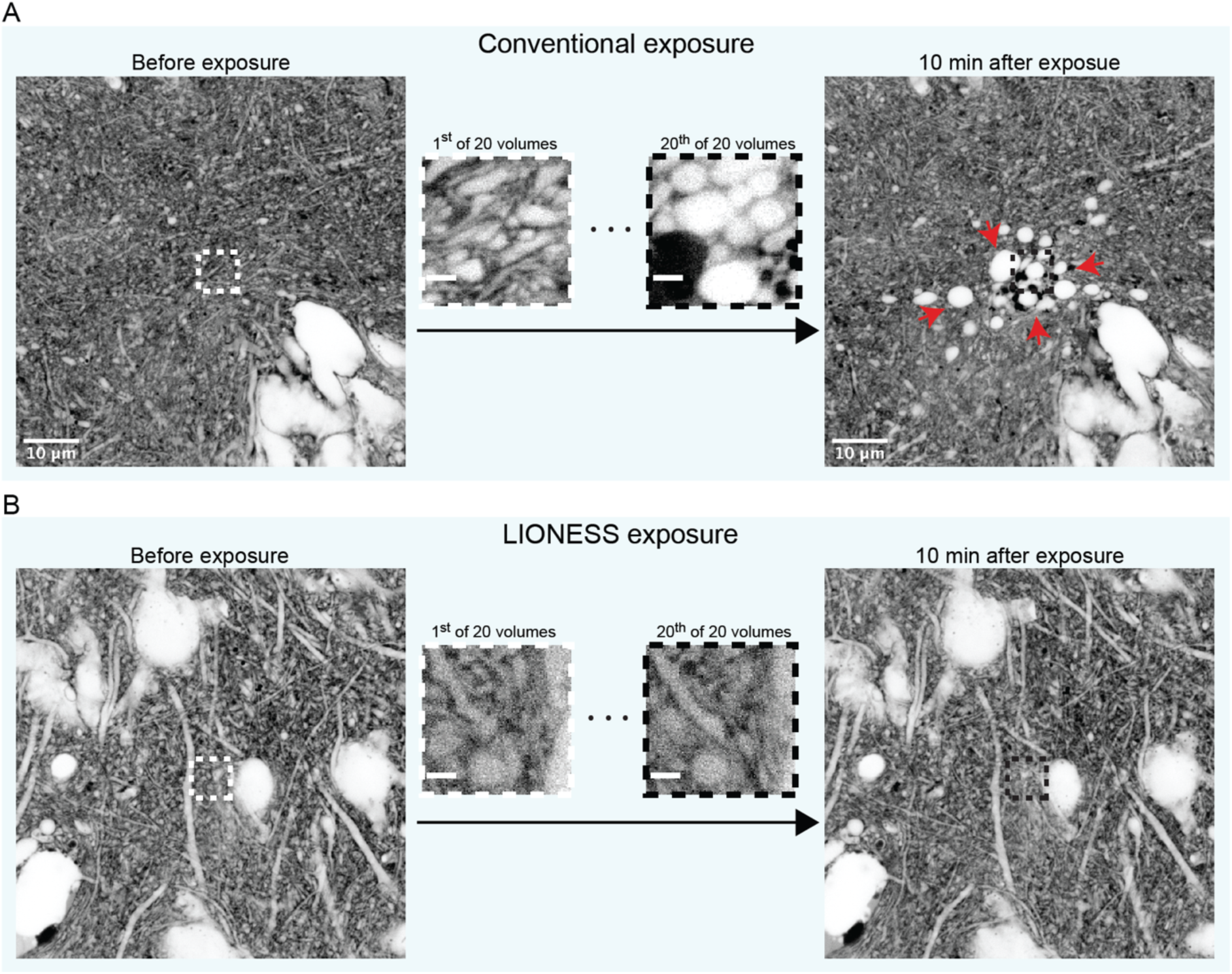
Live-tissue compatibility. (**A**) Confocal overview images in an organotypic hippocampal slice culture before (left) and 10 minutes after (right) scanning a volume in the centre (5 x 5 x 2.5 µm^3^) 20 times in high-photon load STED mode (70 µs voxel integration time). Central images are single planes of the first and last STED volume acquired. Red arrows indicate blebbing and disintegrating cells. (**B**) Confocal overview images of a different region before (left) and 10 minutes after (right) scanning a volume in the centre (5 x 5 x 2.5 µm^3^) 20 times using LIONESS parameters (10 µs voxel integration time). Central images are single planes of the first and last LIONESS volume acquired. Scale bars: confocal: 10 µm, STED: 1 µm.

**Suppl. Fig. 5.**
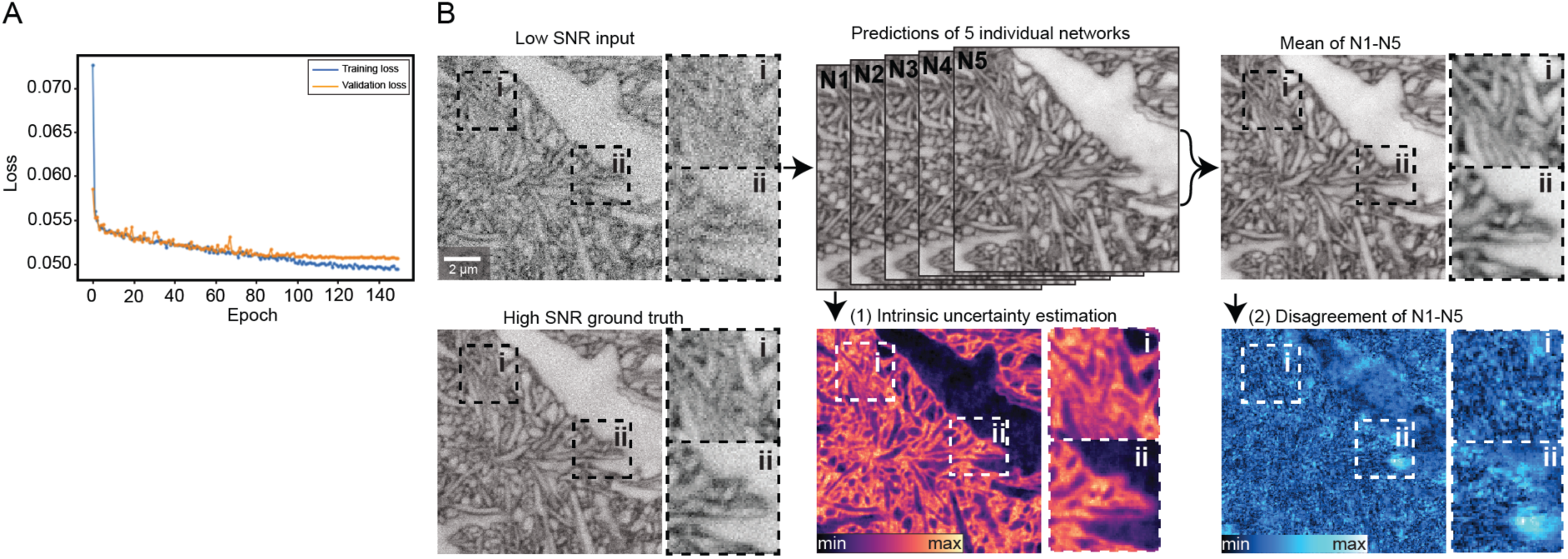
SNR restoration training. (**A**) Training and validation loss as a function of epoch number of the SNR-restoring deep artificial network. (**B**) Validation of artificial network predictions on paired low- and high-SNR data that were not part of the network training, recorded with tissue-optimized STED in extracellularly labelled organotypic hippocampal slice cultures. (1) Intrinsic probabilistic estimation of uncertainty for individual predictions (lower panel, middle) and (2) standard deviation of the mean (disagreement) of 5 trained networks N1-N5 for each voxel (lower panel, right). Raw data and network predictions are maximum intensity projections spanning 150 nm. Scale bar: 2 µm.

**Suppl. Fig. 6.**
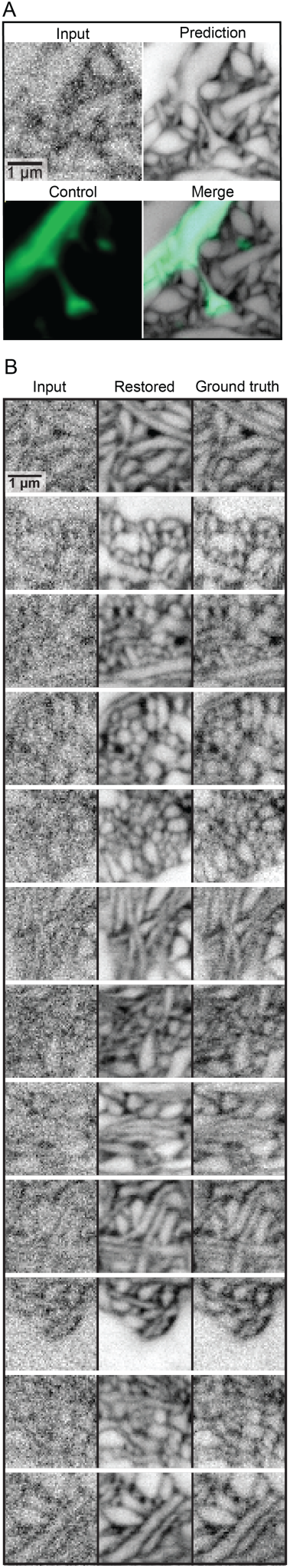
Validation of SNR restoration. (**A**) Raw low-exposure input, SNR-restored artificial neural network prediction, positively labelled (Thy1-EGFP^50^, diffraction limited) control, and overlay of prediction and control in extracellularly labelled organotypic hippocampal slice culture. (**B**) Twelve exemplary areas of raw input, SNR-restored network prediction, and high-SNR ground truth from an imaging volume not included in the network training data. Neuropil in organotypic hippocampal slice culture. Scale bars: 1 µm. Images except for positively labelled control displayed with inverted look-up table. Maximum intensity projections spanning 150 nm. Data used in validation were not part of restoration network training.

**Suppl. Fig. 7.**
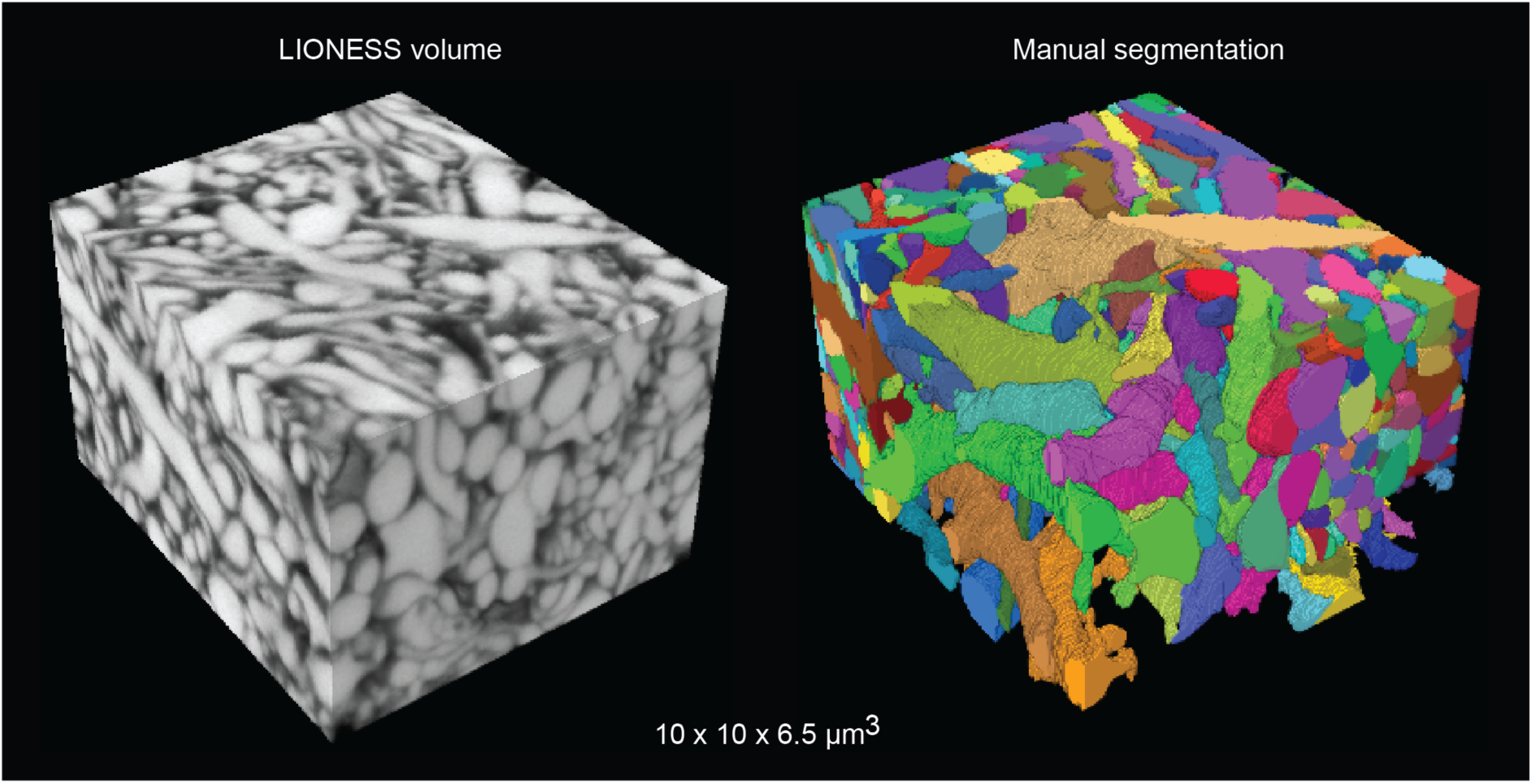
Manual segmentation. *Left*: LIONESS tissue volume in neuropil of organotypic hippocampal slice culture. *Right*: Manual segmentation using VAST Lite 1.3.0 and 1.4.0. The region in the foreground shows a partial segmentation.

**Suppl. Fig. 8.**
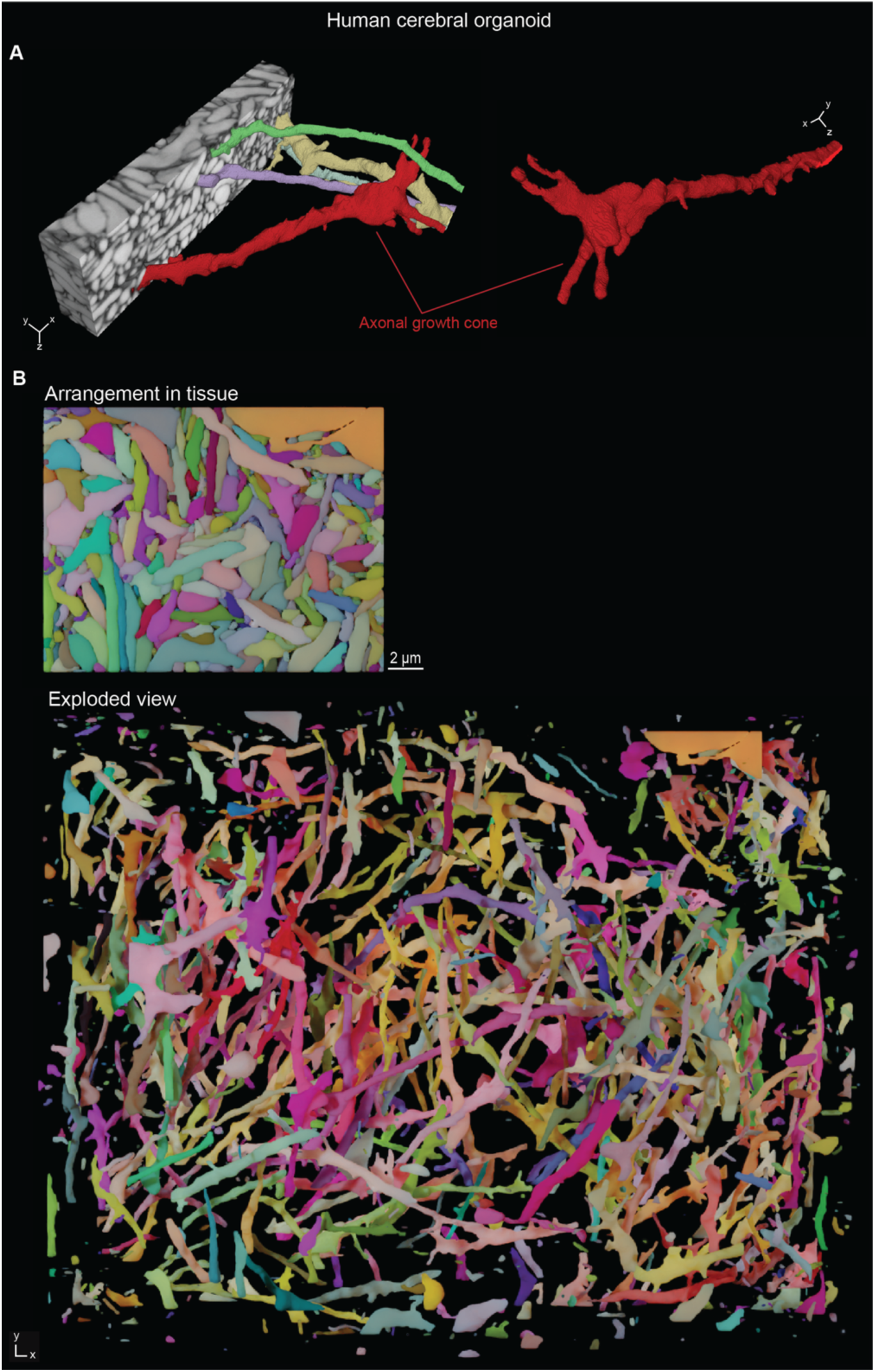
Reconstruction of living human cerebral organoid. (**A**) *Left*: LIONESS volume of the human cerebral organoid in Fig. 1A, eroded to reveal an axonal growth cone transmigrating the dense tissue and a selection of the structures it interacts with. *Right*: The same growth cone viewed from a different angle. (**B**) Top view of the saturated organoid reconstruction in Fig. 1. (Top), and exploded view of the same saturated reconstruction (Bottom). Scale bar: 2 µm.

**Suppl. Fig. 9.**
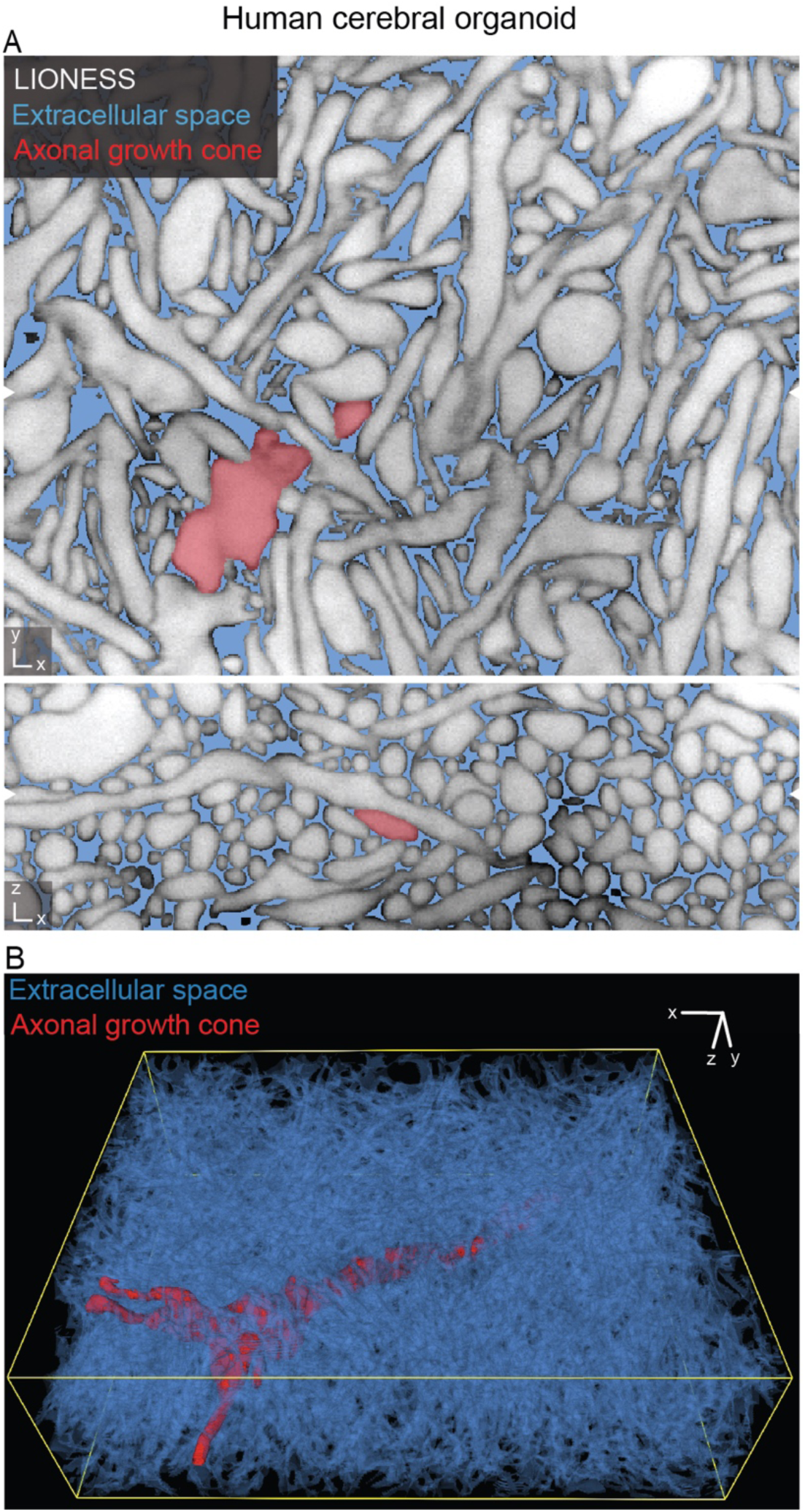
3D-segmentation of extracellular space in a human cerebral organoid. (**A**) Orthogonal views from the human cerebral organoid dataset in Fig. 1A. Extracellular space is highlighted in blue, LIONESS data is shown in grey, and the same axonal growth cone as in Suppl Fig. 8 is indicated in red. White arrowheads at image edges indicate corresponding orthogonal planes. Extracellular space was obtained as the space not occupied by cellular segments. (**B**) 3D-reconstruction of the extracellular space (blue) with the axonal growth cone (red).

**Suppl. Fig. 10.**
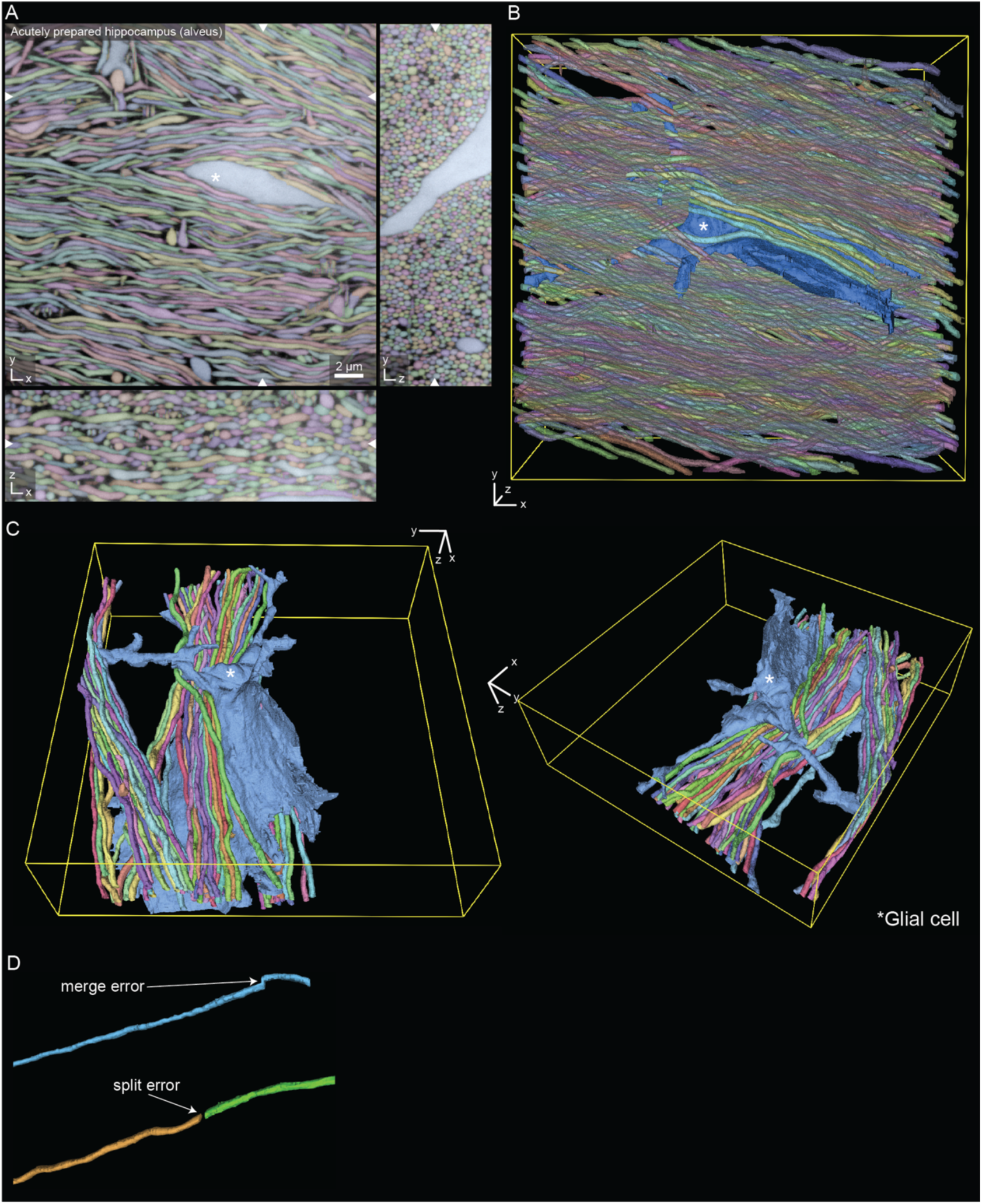
Reconstruction in living hippocampal alveus. (**A**) Three orthogonal planes from a fully segmented LIONESS volume in the alveus region of an acutely prepared mouse hippocampus. The white asterisk indicates a glial cell stretching through dense axons. White arrowheads at image edges indicate the position of orthogonal *xy*-, *yz*- or *xz*-views. Scale bar: 2 µm. (**B**) 3D-rendering of a subset of structures from the same dataset as shown in panel A. (**C**) 3D-reconstruction of the glial cell marked in panel A and selected axons, viewed from two different angles. (**D**) Examples of error types after automated segmentation.

**Suppl. Fig. 11.**
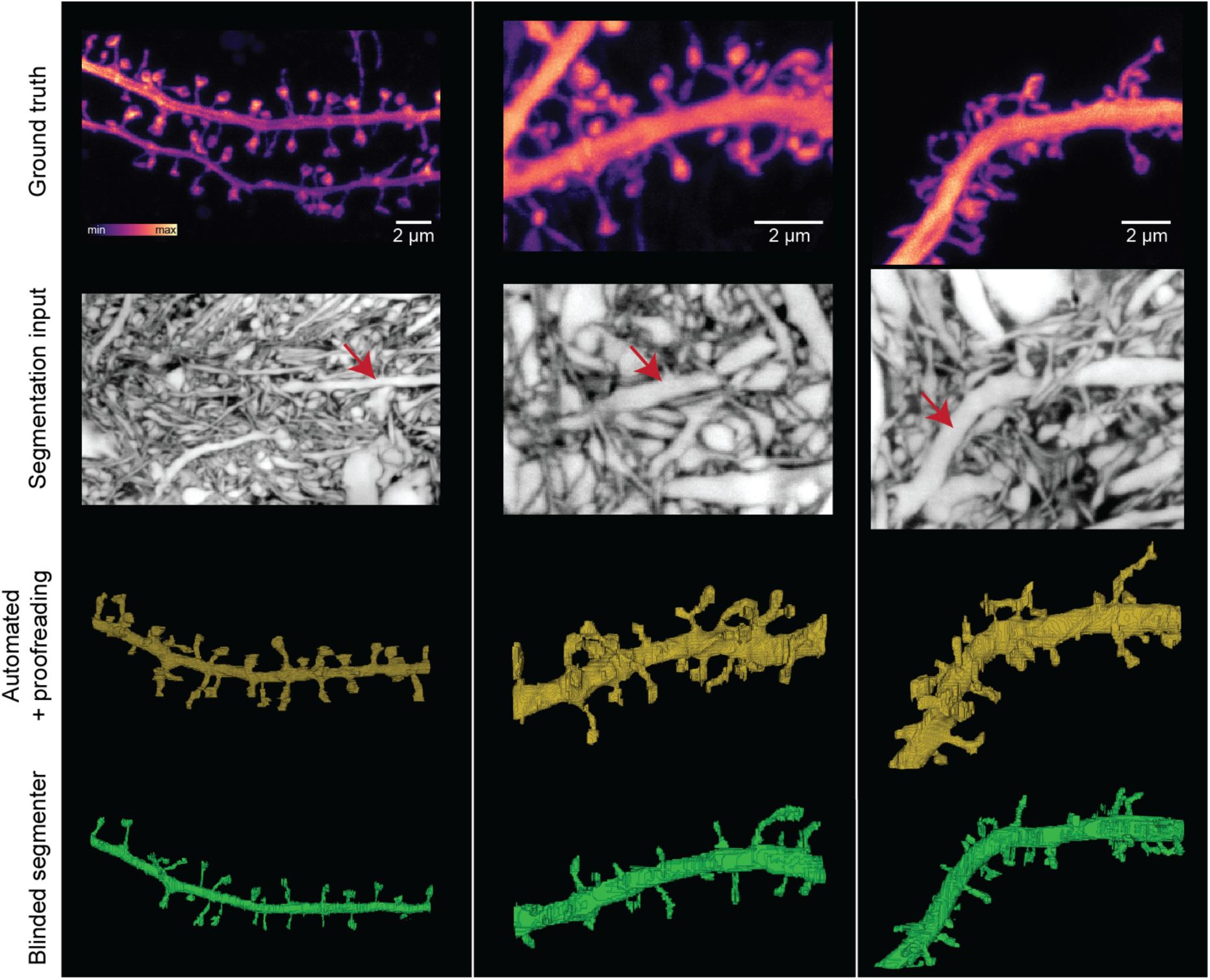
Validation of LIONESS segmentation. *Top row*: Maximum intensity projections of positively labeled (Thy1-EGFP) dendrites from neuropil in 3 different samples of organotypic hippocampal slice cultures, serving as sparse ground truth for LIONESS segmentations. Scale bars: 2 μm. *Second from top*: Volumetric LIONESS acquisitions used as source data for segmentation. Red arrows indicate the dendrites corresponding to the positively labelled structure above. *Third from top*: 3D-reconstructions of LIONESS data with automated segmentation and additional proofreading by the experimenter who recorded the data (i.e. non-blinded to the EGFP channel). *Bottom*: Fully manual spine detection from LIONESS data by a segmenter blinded to the EGFP-channel.

**Suppl. Fig. 12.**
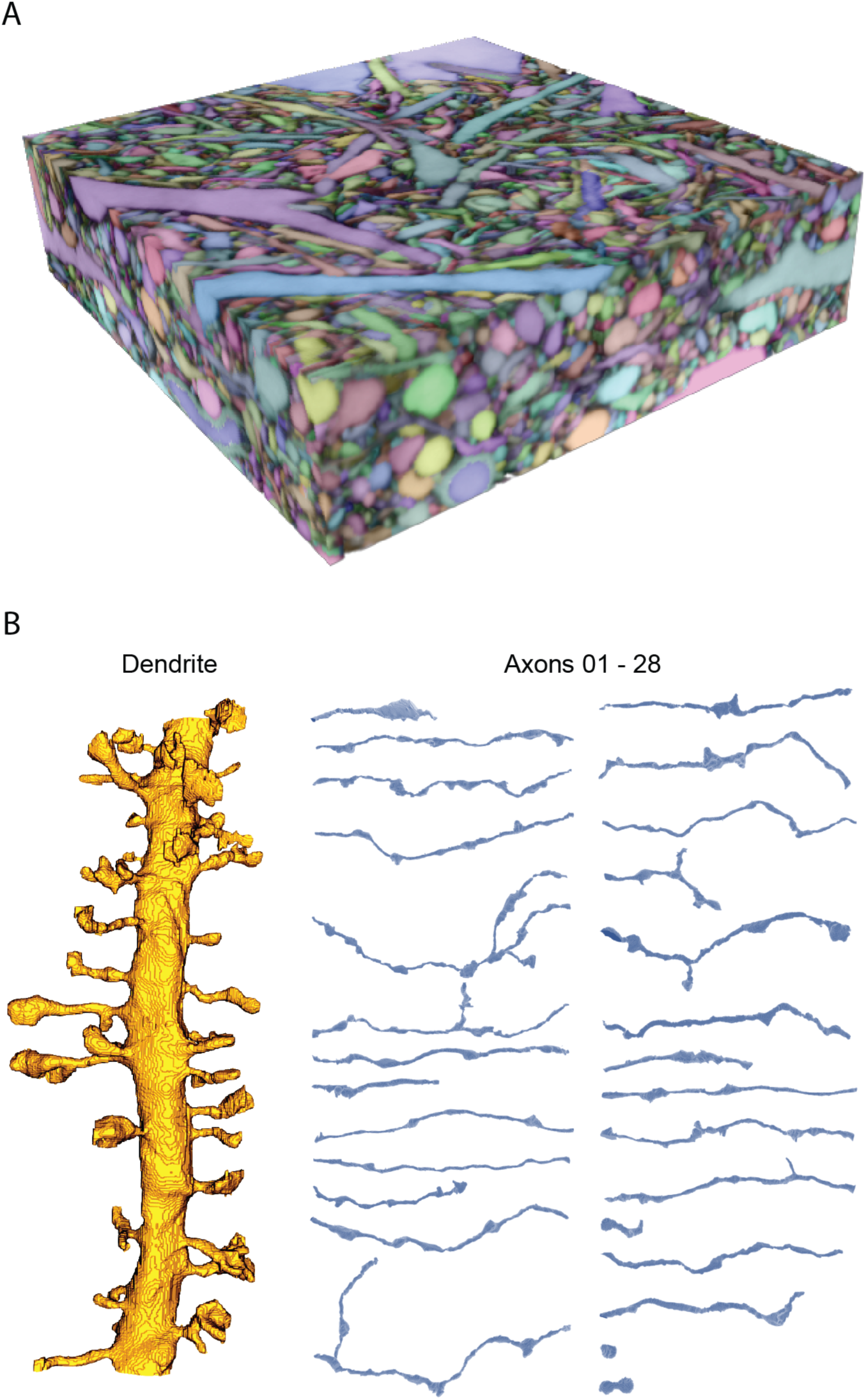
Reconstruction of spiny dendrites and connected axons. (**A**) The entire, automatically segmented dataset from Fig. 3, partially proofread. Volume dimensions: 23.2 x 22 x 6 µm^3^. (**B**) Spiny dendrite and the 28 individual connected axons from Fig. 3 without surface smoothing. The two short segments in the bottom right correspond to one bouton right at the edge of the imaging volume and one bouton which could not be unambiguously assigned to an axon.

**Suppl. Fig. 13.**
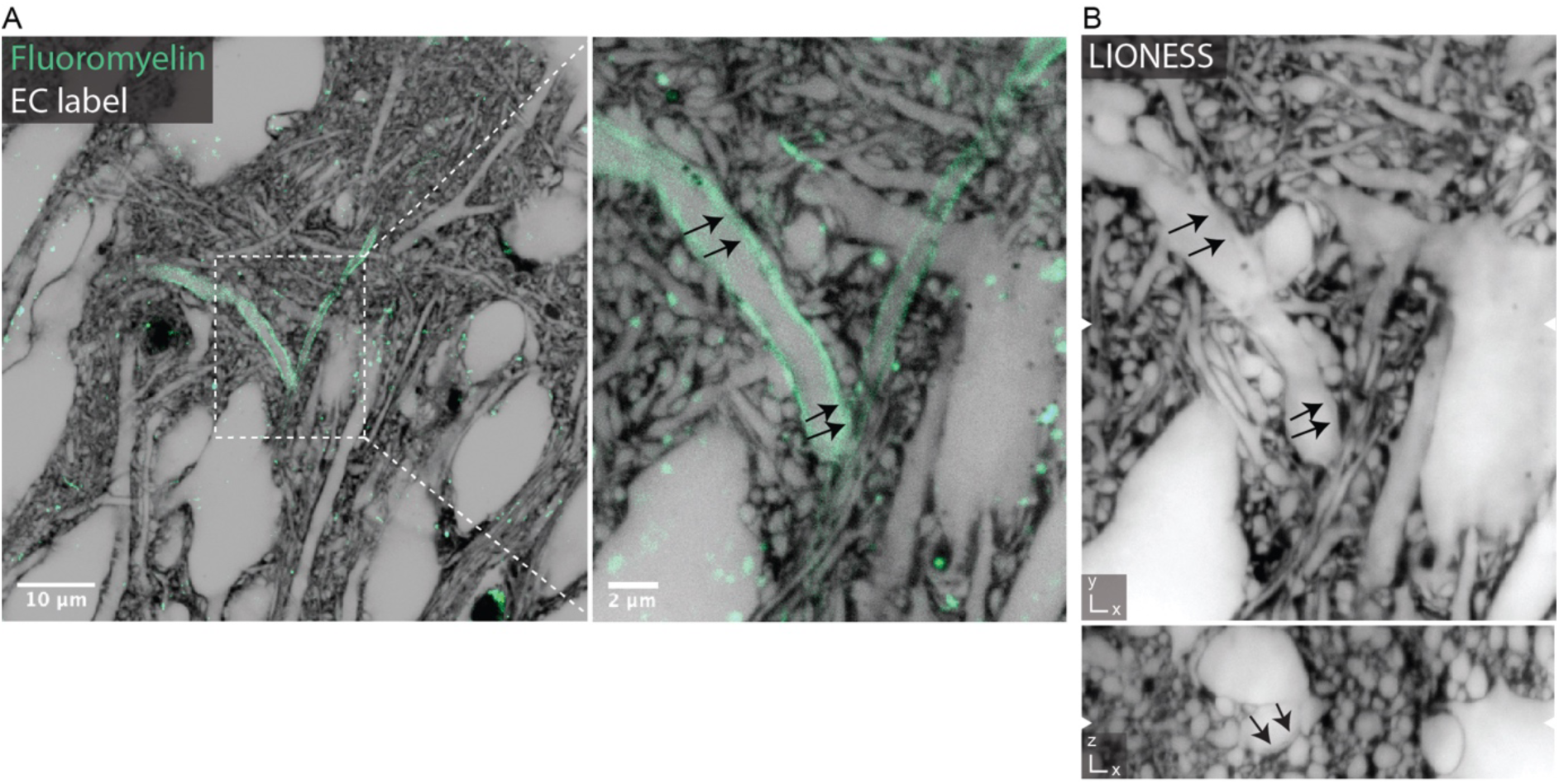
Identification of myelinated axons. (**A**) *Left*: Confocal overview image in organotypic hippocampal slice culture with extracellular (grey, inverted LUT) and additional myelin labelling (Fluoromyelin, green). *Right*: Magnified view highlighting a myelinated axon. Single plane in near-isotropically resolving STED mode for extracellular label and confocal mode for the myelin stain. Scale bars: 10 µm (left), 2 µm (right). (**B**)Volumetric LIONESS acquisition of the same region. Black arrows indicate the border between axon and myelin sheath visible in the LIONESS data. White arrowheads at image edges indicate the corresponding position of *xy*- and *xz*-views. LIONESS images are maximum intensity projections spanning 150 nm.

**Suppl. Fig. 14.**
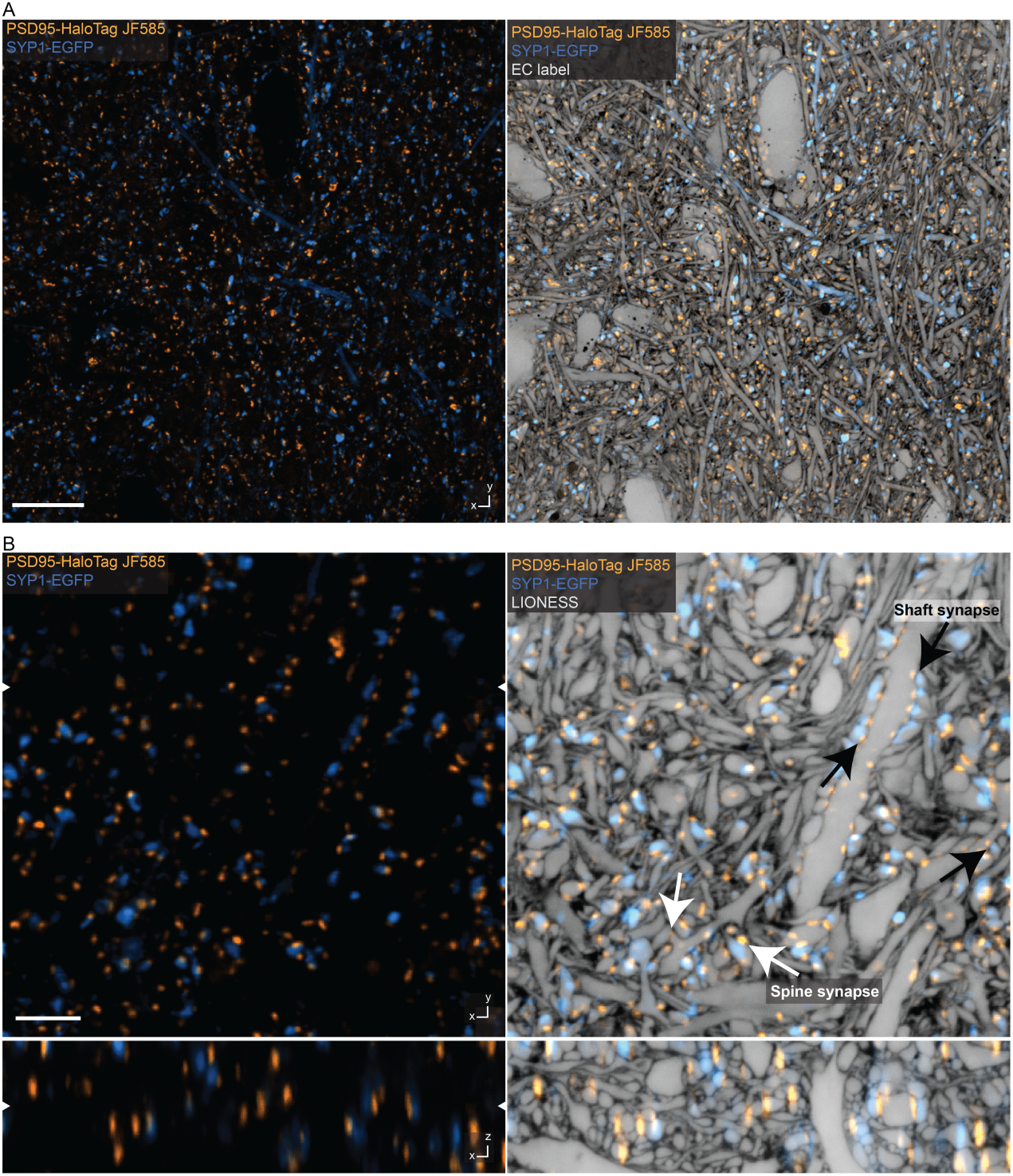
Structural and molecular information. (**A**) Overview image of CA1 hippocampal neuropil in living organotypic slice culture from a transgenic mouse line expressing post synaptic density protein 95 (PSD95)-HaloTag to label excitatory postsynapses (orange, STED). A subset of presynaptic terminals were labelled with a synaptophysin 1 (SYP1)-EGFP fusion protein, encoded by a pseudotyped rabies virus (blue, confocal). *Left*: Molecular markers. HaloTag labelled with JF585. *Right*: Same region with structural context from additional extracellular labeling (STED). Scale bar: 10 µm. (**B**) Orthogonal planes in *xy*- and *xz*-direction of an imaging volume in CA1 from a different sample. Labeling and color coding as in panel A. *Left*: Confocal imaging after denoising^56^. *Right*: Additional overlay with near-isotropically super-resolved LIONESS data, clarifying the relationship of molecularly defined entities (pre- and postsynapses) with cell- and tissue-structure. Diffraction-limited signals extend beyond corresponding structures recorded in LIONESS mode, particularly evident in the *xz*-view. White arrows indicate excitatory spine synapses, black arrows indicate excitatory shaft synapses. White arrowheads at image edges indicate the position of corresponding orthogonal planes (left image). LIONESS images correspond to maximum intensity projections spanning 150 nm. Scale bar: 3 µm.

**Suppl. Fig. 15.**
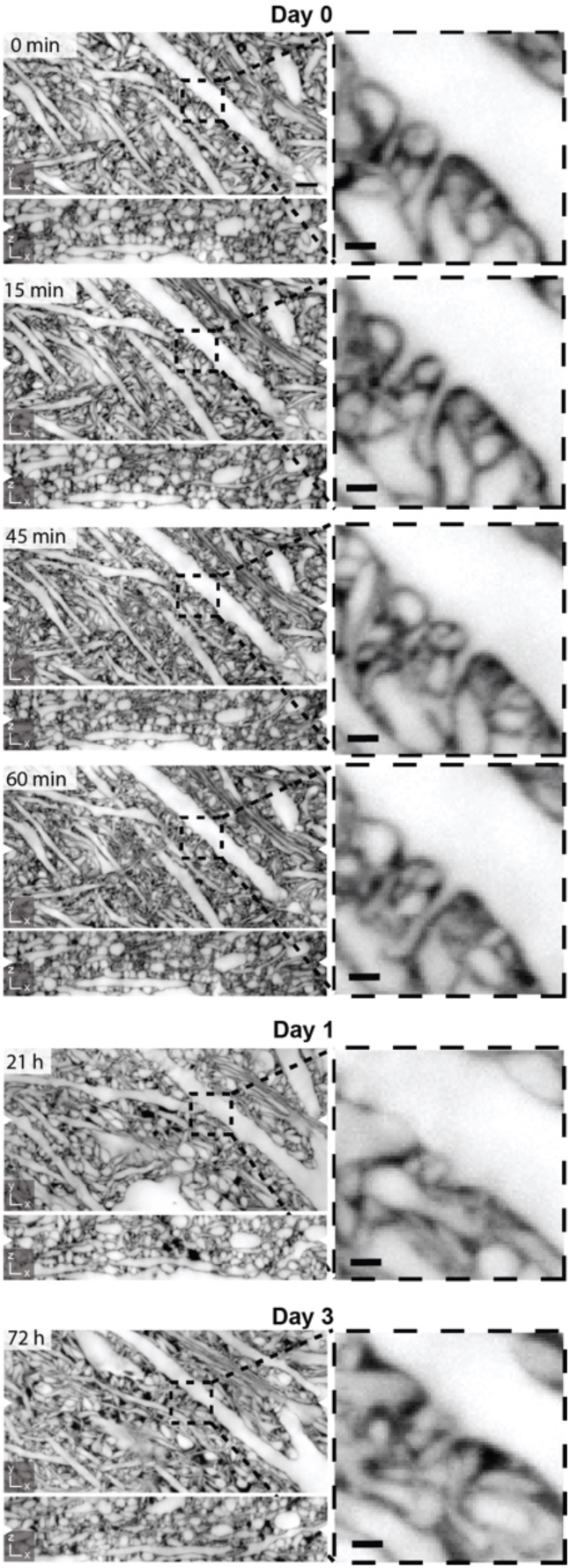
Structural dynamics in repeated volumetric LIONESS acquisition over 3 days. Corresponding orthogonal planes in *xy*- and *xz*-direction from 6 consecutive LIONESS measurements of the same volume in the neuropil of an organotypic hippocampal slice culture. The volume was initially imaged 4 times within one hour and then again after one day and after three days. This indicates tissue viability after repeated volumetric LIONESS imaging. Magnified views: Subregion with dendritic spines revealing morphodynamics. Scale bars, overview: 2 µm, magnified views: 500 nm. White arrowheads at image edges indicate the position of corresponding orthogonal planes. Maximum intensity projections spanning 150 nm. Additional dark regions on day 1 and day 3 likely represent branched processes of a damaged cell that took up dye after repeated manual mounting of the sample (supported by a membrane for interface tissue culture), transfer to the microscope, volumetric imaging, unmounting, and transfer back to the tissue culture incubator.

**Suppl. Fig. 16.**
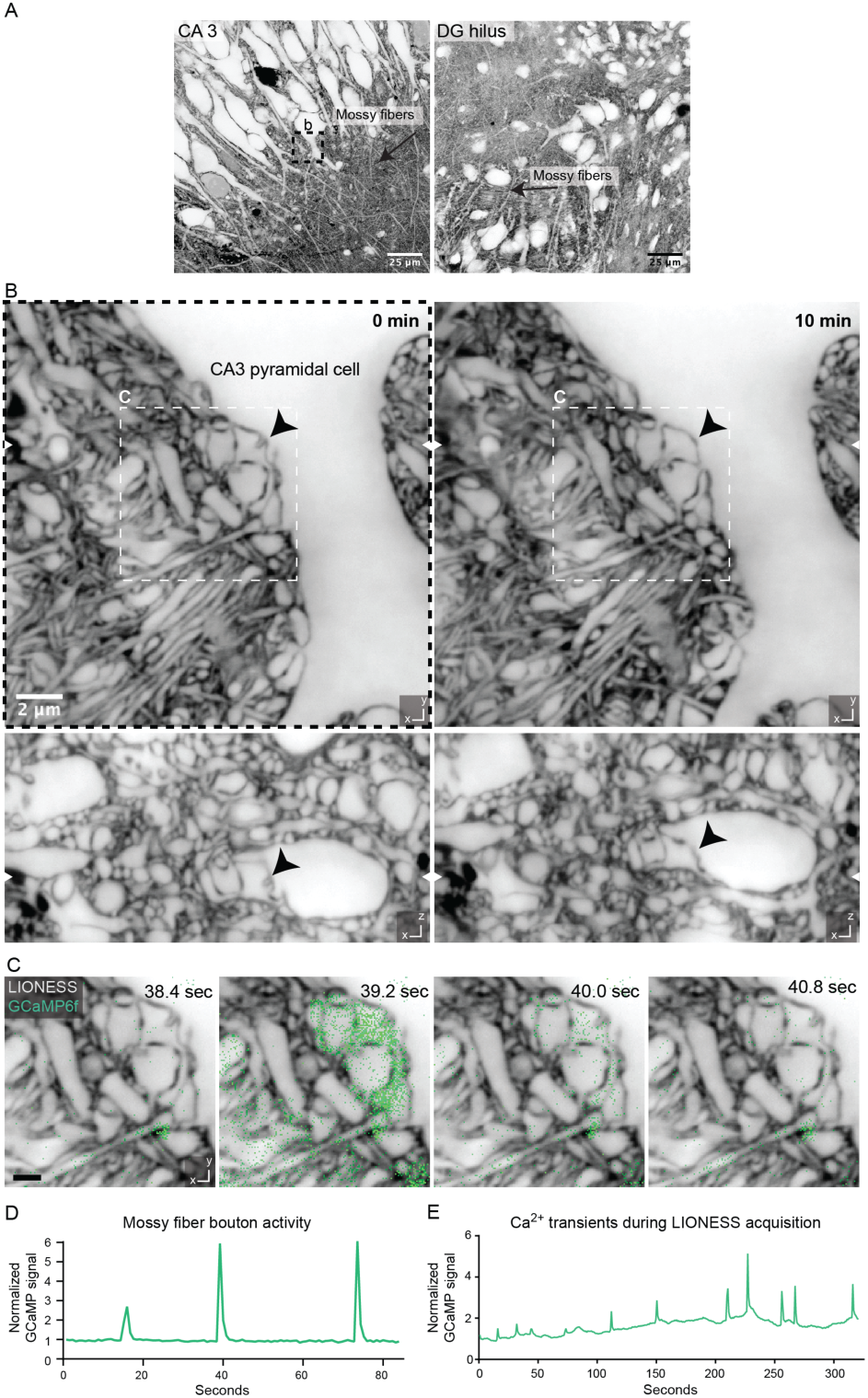
Correlating structure and morphodynamics with Ca^2+^-activity. (**A**) Overview images in organotypic hippocampal slice culture with mossy fibers conveying excitatory input from DG granule cells (right) to CA3 pyramidal neurons (left). Scale bar: 25 µm. (**B**) Volumetric LIONESS acquisitions in the stratum lucidum of CA3 at two timepoints (left: 0 minutes, right: 10 minutes) revealed morphodynamics of the complex interface between pre- and post-synaptic structures at mossy fiber to CA3 pyramidal neuron synapses. The black arrowhead marks a structure changing over time. White arrowheads at image edges indicate the corresponding positions of *xy-* and *xz-* views. Scale bar: 2 µm. (**C**) Plane from the LIONESS volume overlaid with diffraction-limited signal from the calcium indicator GCaMP6f (green). LIONESS images are identical replicates providing structural context to the time-varying Ca^2+^-signals. Scale bar: 1 µm. (**D**) GCaMP signal of the mossy fiber bouton shown in (C) as a function of time. Total signal from a rectangular region enclosing the mossy fiber bouton was integrated and normalized to the first frame. (**E**) GCaMP signal as a function of time (and position) recorded as an additional color channel during the volumetric LIONESS acquisition for timepoint 0 min in (B), indicating that Ca^2+^-activity continued during LIONESS acquisition. LIONESS images are maximum intensity projections spanning 150 nm.

**Suppl. Fig. 17.**
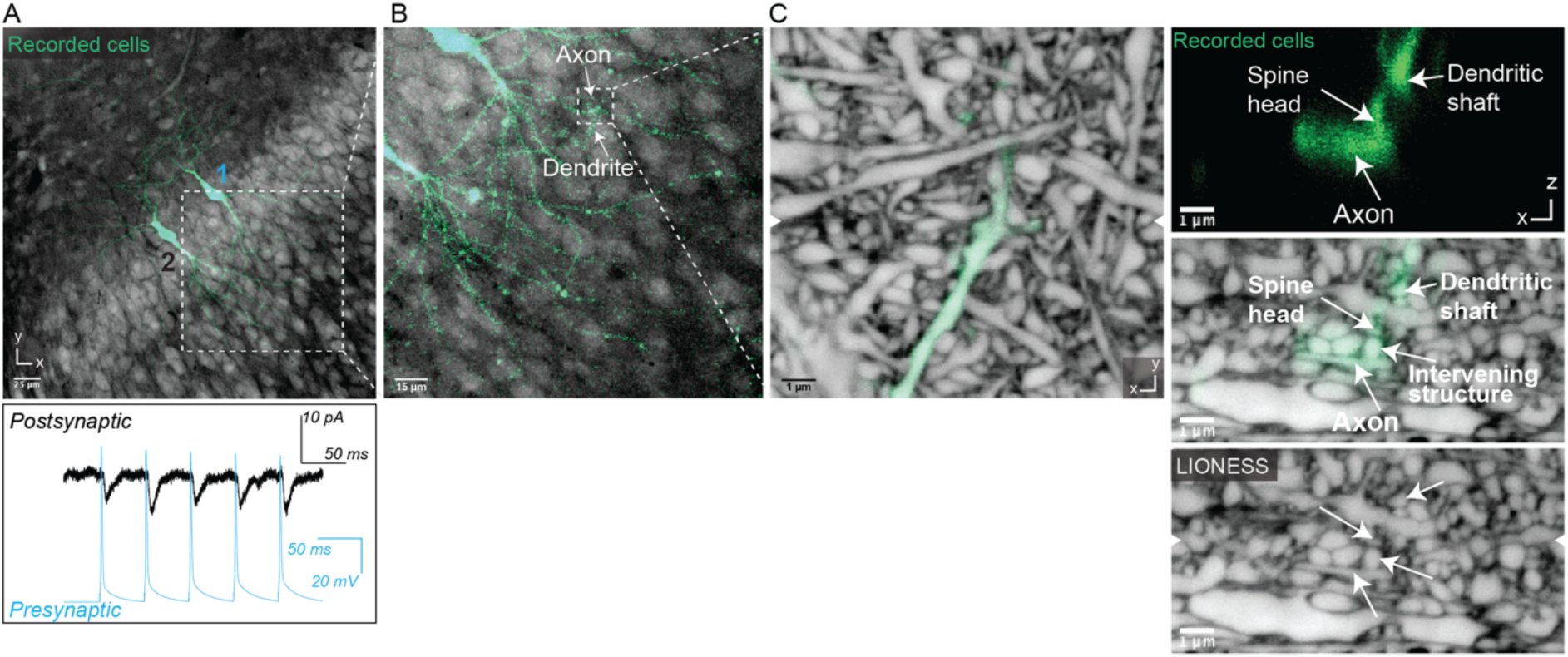
Correlating structural with electrophysiological information. (**A**) Two CA1 pyramidal neurons after patch clamp recording and filling with fluorescent dye in living organotypic hippocampal slice culture. Currents in neuron (2, black trace in bottom panel) elicited by triggered action potentials in neuron (1, blue trace in bottom panel) reveal a monosynaptic connection in the paired recording. Scale bar: 25 µm. Confocal image of positively labelled neurons (green) and extracellular label (grey) with low-numerical aperture objective. (**B**) Region where axon of neuron 1 overlaps with a dendrite of neuron 2, suggesting a synaptic connection in confocal imaging. Scale bar: 15 µm. (**C**) Detailed view of overlap region with positively labelled structures (green) read out at diffraction-limited resolution with a high-numerical aperture objective, embedded in surrounding volume recorded with LIONESS. Orthogonal views in *xy*- and *xz*-directions, with arrowheads at image edges indicating the position of the corresponding orthogonal sections. The diffraction-limited *xz*-view of the positively labelled structures indicated a synaptic connection (top). The increased resolution and comprehensive labelling of all cellular structures in the LIONESS xz-view disclosed an intervening structure unrelated to the two patch-clamped neurons. Scale bars: 1 µm. Maximum intensity projections spanning 150 nm.

**Suppl. Fig. 18.**
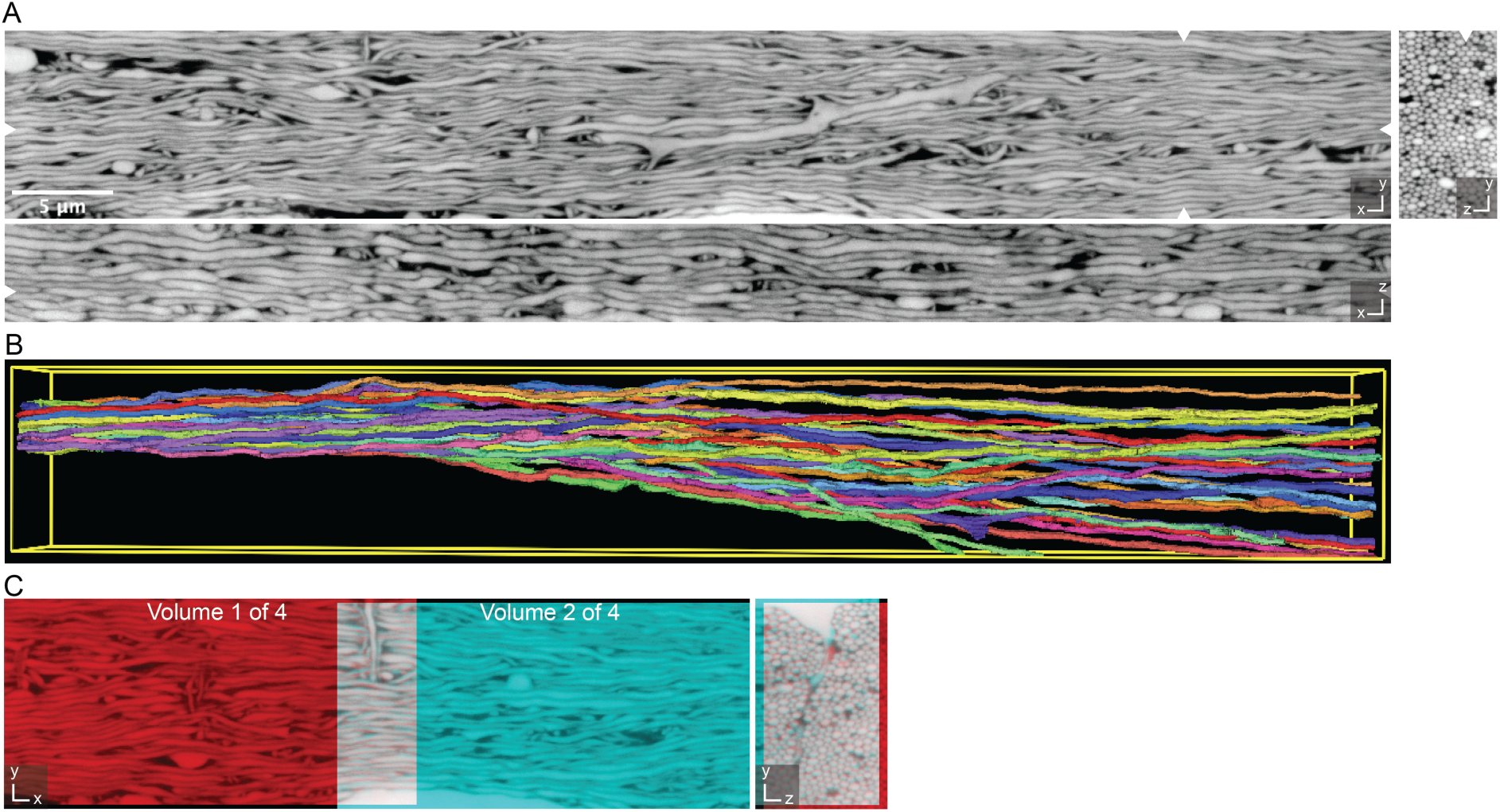
Extending LIONESS tissue volumes. (**A**) Orthogonal planes in *xy*-, *xz*-, and *yz*-directions from a LIONESS volume in the alveus region of an acutely prepared mouse hippocampus. Data were registered from 4 consecutive, partially overlapping acquisitions. White arrowheads at image edges indicate position of corresponding orthogonal planes. Maximum intensity projections spanning 150 nm. Scale bar: 5 µm. (**B**) 3D-rendering of selected axons from (A), forming a tight bundle in the left and progressively fanning out. (**C**) Example of alignment between two of the partially overlapping subvolumes in *xy*- and *yz*-views. Individual subvolumes are shown in red and cyan, such that overlapping regions add up to white color, indicating the degree of overlap.

**Suppl. Fig. 19.**
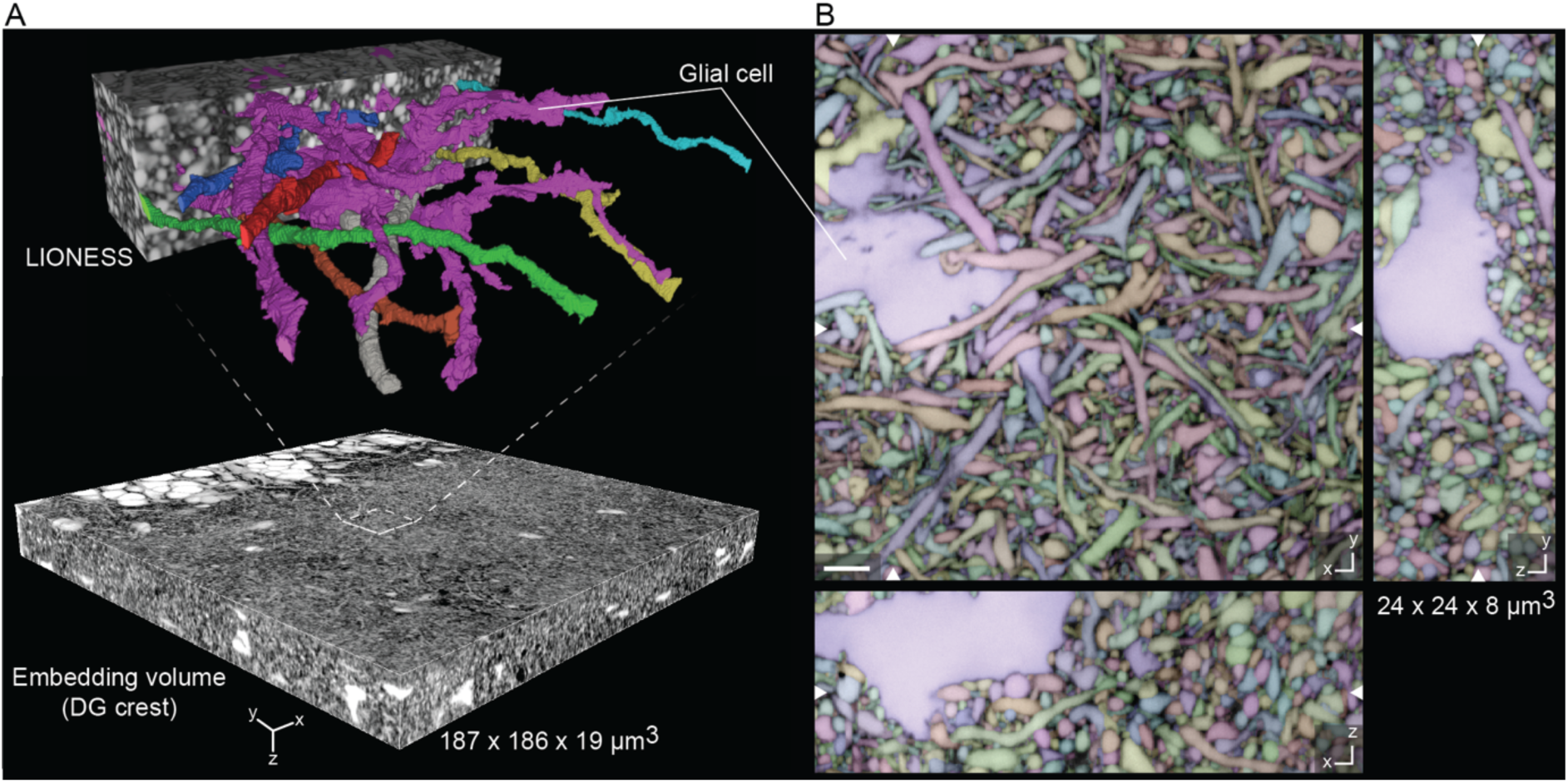
Meso-scale tissue context. (**A**) Meso-scale overview volume acquired in diffraction-limited mode with select subvolume acquired and reconstructed using LIONESS in an organotypic hippocampal slice culture. A glial cell is 3D-rendered together with exemplary neuronal processes, showing their mutual arrangement. (**B**) Three orthogonal planes from the automated segmentation of the LIONESS volume in (A). Segmentation (color) and LIONESS data are overlaid. No proofreading was applied. White arrowheads at image edges indicate corresponding orthogonal planes. The same glial cell is indicated in both panels. Scale bar: 2 µm.

**Suppl. Fig. 20.**
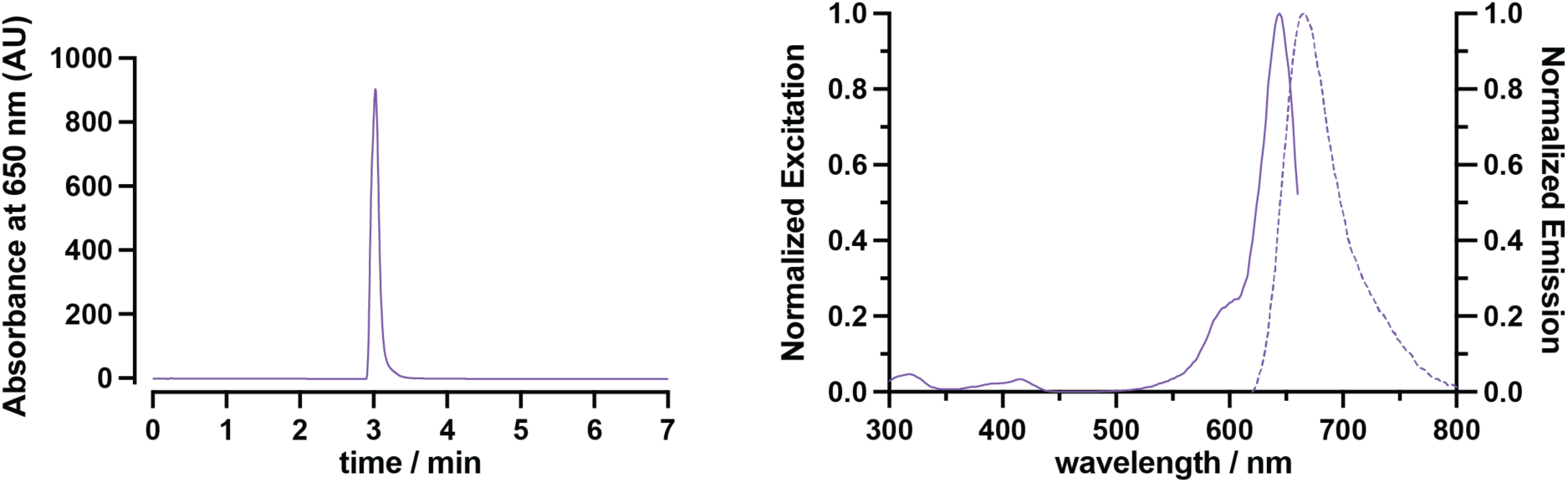
Characterization of SulfoAtto 643. *Left*: Absorbance as a function of elution time acquired by liquid chromatography low resolution mass spectrometry. *Right*: Normalized excitation and emission spectra of SulfoAtto 643.

**Supplementary Video 1.**

**LIONESS volume in living hippocampal alveus - 1 (*xy* view).** Full LIONESS stack (*xy*-view) in the alveus region of an acutely prepared mouse hippocampus, corresponding to the dataset in Suppl. Fig. 10. A glial cell embedded in dense axonal structures oriented in various direction can be appreciated. Step size: 50 nm (160 optical sections).

**Supplementary Video 2.**

**LIONESS volume in living hippocampal alveus - 2 (*yz* view).** Full LIONESS stack (*yz*-view) in the alveus region of an acutely prepared mouse hippocampus, corresponding to the dataset in Suppl. Fig. 10. A glial cell embedded in dense axonal structures can be appreciated. Of note, axons are organized in layers with different main orientation, apparent from the differential movement of axonal cross sections in the fly through. Step size: 50 nm (500 optical sections).

**Supplementary Video 3.**

**LIONESS volume in living hippocampal neuropil.** Full LIONESS stack (x*y*-view, followed by *xz*-view) of a volume of dentate gyrus in organotypic hippocampal slice culture, corresponding to the dataset in Fig. 3. Various neuronal and non-neuronal structures are visible. Step size: 50 nm (xy-view: 120 optical sections, xz-view: 440 optical sections).

**Supplementary Video 4.**

**Molecularly informed LIONESS volume in living hippocampal neuropil.** Full LIONESS stack (*xy*-view) of the entire PSD95 (orange) and synaptophysin (blue) labelled dataset shown cropped in Fig. 4. Molecular information confirms synapse location in relation to the structural LIONESS measurement. Step size: 50 nm (120 optical sections).

**Supplementary Video 5.**

**GCaMP recording, overview.** Diffraction limited recording of calcium transients using Ai95/Prox1-cre mouse organotypic hippocampal slice cultures extracellularly labelled with Atto 643. The time series corresponds to a single plane of a region in the DG during application of the GABA_A_ receptor antagonist gabazine. Acquisition frame rate was 1.25 Hz.

**Supplementary Video 6.**

**GCaMP recording, single synapse.** Full time series of the GCaMP recording after gabazine application in Ai95/Prox1-cre mouse organotypic hippocampal slice cultures shown in Suppl. Fig. 16C. Ca^2+^ transients of a mossy fiber bouton attached to a CA3 pyramidal neuron are visible. LIONESS images are identical replicates providing structural context to the time-varying Ca^2+^-signals (green). Acquisition frame rate of the GCaMP signal was 1.25 Hz.

**Supplementary Video 7.**

**GCaMP recording, chemogenetically activated single synapse.** Full series of the GCaMP recording shown in Fig. 5A. Ca^2+^ transients of a mossy fiber bouton attached to a hilar mossy cell in Ai95/Prox1-cre mouse organotypic hippocampal slice cultures are visible. LIONESS and dTomato (orange, coexpressed with the DREADD hM3Dq) images are identical replicates placing the overlaid time-varying Ca^2+^-signals (green) after stimulation with the DREADD ligand CNO into structural context. Acquisition frame rate of the GCaMP signal was 2 Hz.

**Supplementary Video 8.**

**3D reconstruction of mossy fiber bouton on hilar mossy cell, including neighboring boutons.** Same reconstruction as in Fig. 5B, but including additional, hM3Dq-negative mossy fiber boutons from automated segmentation (left), and the thorny excrescences alone (right). Additional boutons are not proof-read.

**Supplementary Video 9.**

**LIONESS volume in living hippocampal dentate gyrus - 1 (*xy* view).** Full LIONESS stack (x*y*-view) of a volume of dentate gyrus in organotypic hippocampal slice culture, corresponding to the dataset in Suppl. Fig. 19. Step size: 50 nm (160 optical sections).

**Supplementary Video 10.**

**LIONESS volume in living hippocampal dentate gyrus - 2 (*xz* view).** Full LIONESS stack (x*z*-view) of a volume of dentate gyrus in organotypic hippocampal slice culture, corresponding to the dataset in Suppl. Fig. 19. Step size: 50 nm (480 optical sections).

